# Inflammatory blockade prevents injury to the developing pulmonary gas exchange surface in preterm primates

**DOI:** 10.1101/2021.06.24.449766

**Authors:** Andrea Toth, Shelby Steinmeyer, Paranthaman Kannan, Jerilyn Gray, Courtney M. Jackson, Shibabrata Mukherjee, Martin Demmert, Joshua R. Sheak, Daniel Benson, Joe Kitzmiller, Joseph A. Wayman, Pietro Presicce, Christopher Cates, Rhea Rubin, Kashish Chetal, Yina Du, Yifei Miao, Mingxia Gu, Minzhe Guo, Vladimir V. Kalinichenko, Suhas G. Kallapur, Emily R. Miraldi, Yan Xu, Daniel Swarr, Ian Lewkowich, Nathan Salomonis, Lisa Miller, Jennifer S. Sucre, Jeffrey A. Whitsett, Claire A. Chougnet, Alan H. Jobe, Hitesh Deshmukh, William J. Zacharias

## Abstract

Malformations of or injuries to the developing lung are associated with perinatal morbidity and mortality with lifelong consequences for subsequent pulmonary health. One fetal exposure linked with poor health outcomes is chorioamnionitis, which impacts up to 25-40% of preterm births. Severe chorioamnionitis with prematurity is associated with significantly increased risk of pulmonary disease and secondary infections in childhood, suggesting that fetal inflammation may significantly alter developmental ontogeny of the lung. To test this hypothesis, we used intra-amniotic lipopolysaccharide (LPS, endotoxin) to generate experimental chorioamnionitis in prenatal Rhesus macaque (*Macaca mulatta*), a model which shares critical structural and temporal aspects of human lung development. Inflammatory injury directly disrupts the developing gas exchange surface of the primate lung, with extensive damage to alveolar structure, particularly the close association and coordinated differentiation of alveolar type 1 pneumocytes and specialized alveolar capillary endothelium. Single cell RNA sequencing analysis defined a multicellular alveolar signaling niche driving alveologenesis which was extensively disrupted by perinatal inflammation, leading to loss of gas exchange surface and alveolar simplification similar to that found in chronic lung disease of newborns. Blockade of IL1β and TNFα ameliorated endotoxin-induced inflammatory lung injury by blunting stromal response to inflammation and modulating innate immune activation in myeloid cells, restoring structural integrity and key signaling networks in the developing alveolus. These data provide new insight into the pathophysiology of developmental lung injury and suggest that modulating inflammation is a promising therapeutic approach to prevent fetal consequences of chorioamnionitis.

## Introduction

The development of a functional gas exchange surface in the fetal lung is required for survival after birth. Lung development in all mammals begins with budding of the lung from the anterior foregut endoderm, followed by stereotypic branching of lung tubules that generates the complex airway tree^1–3^. Proliferation and differentiation of epithelial cells in the distal lung generates alveoli, which make up the gas exchange surface of the mature lung^4^. In humans and other primates, cellular differentiation and alveolar formation begins *in utero*, during the third trimester of pregnancy, and continues until adolescence^5–7^. Alveolar type 2 (AT2) cells generate surfactant proteins and lipids to modulate surface tension in the alveoli^8^, and alveolar type 1 (AT1) cells associate closely with alveolar endothelial cells to form a large surface for exchange of oxygen and carbon dioxide^4,9,10^. Together, these processes are termed alveologenesis. At birth, fluid is reabsorbed and cleared from the alveolar space, and ventilation begins with air entry into the alveolus, accompanied by reduction of pulmonary vascular resistance, increasing capillary blood flow to match ventilation, allowing adequate oxygenation at birth^11,12^. When these carefully coordinated systems fail, neonatal respiratory distress results, which can be rapidly fatal without neonatal intensive care.

While cellular and genetic mechanisms directing lung development have been elucidated by studies of murine lung, there are important differences in structural and temporal development between mouse models and human lungs which limit applicability of murine studies to human disease^13,14^. First, recent studies demonstrate important differences in the cell-specific expression of transcription factors (TFs) and signaling molecules in humans compared to mice, both during development^15,16^ and in adulthood^14,17^. Second, while rodents have a discrete bronchial-alveolar duct junction (BADJ) (Figure S1C), humans and other primates have a distinct secretory epithelium lining respiratory bronchioles and alveolar ducts^18^, common sites of damage in pulmonary diseases (Figure S1A-B)^19^. Finally, while murine alveologenesis occurs between postnatal day 3 (PND3) and PND14, human alveologenesis begins *in utero*^5–7^.

Defects in lung morphogenesis and lack of pulmonary surfactant are common causes of respiratory failure in the perinatal period, requiring intensive care to support ventilation^8^. Incomplete AT2 cell differentiation causes surfactant deficiency and respiratory distress syndrome (RDS)^20^, marked by ventilatory failure and hypoxemia requiring respiratory support. While antenatal glucocorticoids and surfactant replacement therapies have improved morbidity and mortality from RDS^21–23^, premature infants who require intensive care for RDS and other pulmonary pathologies may develop chronic lung disease of prematurity (CLDP) or bronchopulmonary dysplasia (BPD)^24^. These disorders are characterized by alveolar simplification, reduced gas exchange surface, and the need for long-term ventilatory support or supplemental oxygen.

The common perinatal stressor chorioamnionitis, defined as inflammation of the chorion and/or the amnion of the placenta, complicates as many as 25-40% of preterm births^25–27^. Importantly, chorioamnionitis is associated with increased risk of multiple respiratory comorbidities following delivery^28–30^, and is considered a distinct risk factor associated with the development of respiratory disease in childhood and later life^31^. Extensive prior literature has demonstrated that chorioamnionitis is associated with increased pro-inflammatory cytokines in the amniotic fluid, which bathes the developing lung in utero^32,33^. The inflammatory milieu of chorioamnionitis is hypothesized to cause tissue injury and remodeling leading to lung disease in survivors.

We sought to investigate how inflammatory injury to the developing fetus alters the trajectory of lung development in a clinically relevant and developmentally appropriate non-human primate Rhesus macaque (*Macaca mullata*) model. Using a combination of histological analysis, high content confocal imaging, and single cell RNA sequencing, we show that Rhesus lung development recapitulates critical aspects of human alveologenesis in the third trimester of gestation, with serial differentiation of epithelial, endothelial, and mesenchymal lineages required for morphological and physiological maturation of the distal lung. Developmental inflammation caused by intra-amniotic endotoxin leads to extensive lung injury, with significant disruption of alveolar structure, effacement of septal structures, injury to progenitor lineages throughout the alveolus, and loss of gas exchange surface. Endotoxin directly damages cell lineages and alters developmental signaling between alveolar cells, with substantial changes to signaling in both mesenchymal cells and lung innate immune lineages. In combination, these insults lead to alveolar simplification histologically similar to CLDP in human neonates. Blockade of the major chorioamnionitis-associated cytokines IL1β and TNFα protects the developing lung from inflammatory injury, preventing histological disruption and restoring the cellular quorum and signaling milieu of the alveolus. Together, these data provide new insight into key mechanisms of developmental lung injury and highlight targeted inflammatory blockade as a potential therapeutic approach to ameliorate lung injury in the vulnerable neonatal population.

## Results

### Inflammatory injury disrupts the developing alveolar gas exchange surface

To evaluate the dynamics of the development of the gas exchange surface in late gestation primates, we examined Rhesus lung at multiple late prenatal time points (Figure 1A-J, S1D-F, S2A-O; extensive additional histological and IHC images of all ages of Rhesus lung development are available online at https://research.cchmc.org/lungimage/).

**Figure 1.**
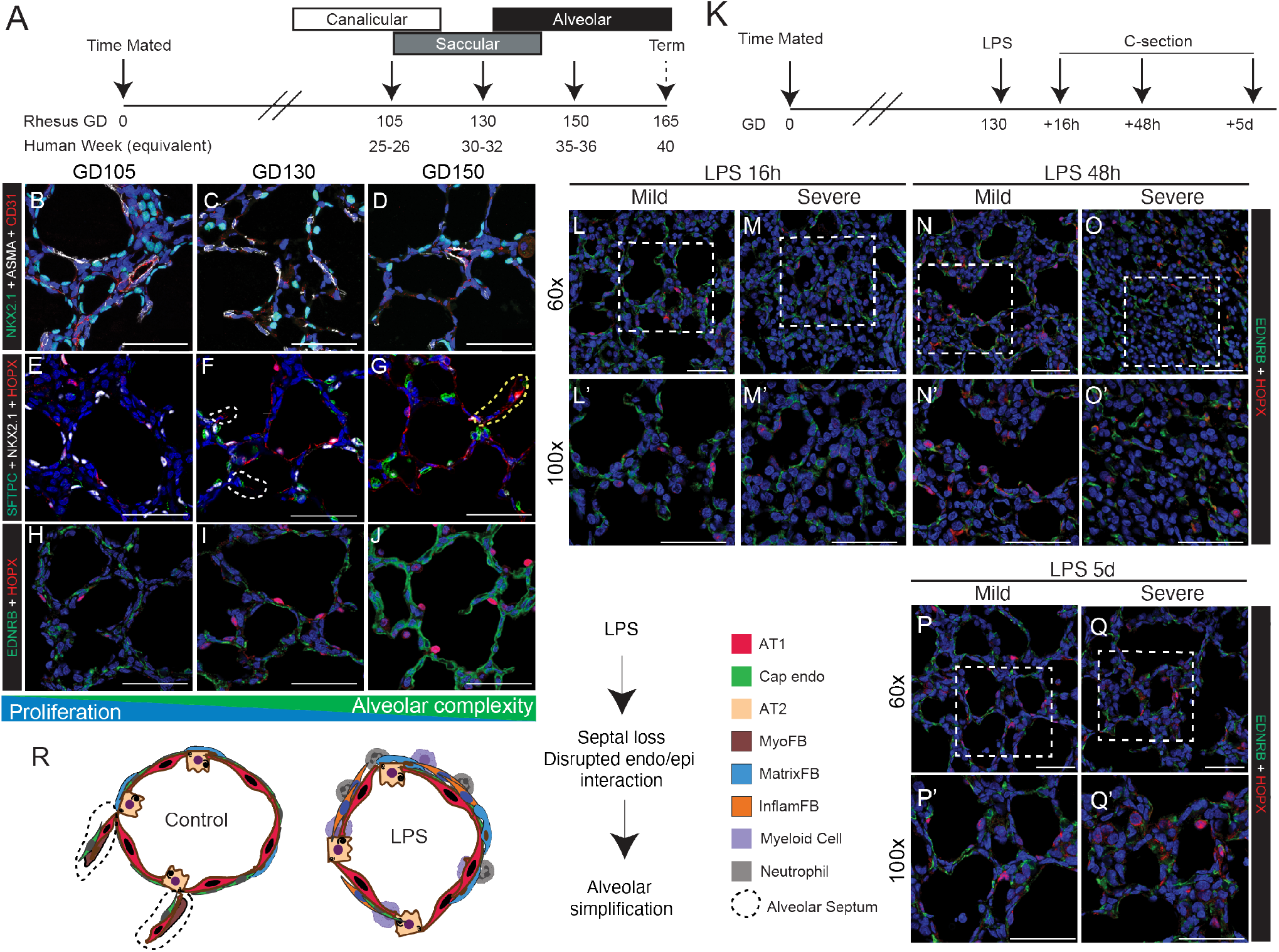
Alveolar simplification and loss of gas exchange surface after perinatal inflammatory injury. (A) Schematic of experimental design for controls, with planned harvest at gestational day (GD) timepoints representative of each stage of lung maturation (GD105 [canalicular; n=3], GD130 [saccular; n=11], GD150 [alveolar; n=3]). (B-J) Confocal microscopy of the developing gas exchange surface highlighting epithelial interactions with underlying mesenchyme and endothelium. (B-D) NKX2.1^+^ epithelial cells (green) form close interactions with both ASMA^+^ myofibroblasts (white) and CD31^+^ endothelial cells (red). (E-G) Epithelial differentiation of HOPX^+^ AT1 cells (red) and SFTPC^+^ AT2 cells (green) increase as development progresses. (H-J) Interactions of HOPX^+^ AT1 cells (red) and EDNRB^+^ alveolar capillary endothelial cells (aerocytes; green) increase during patterning of the developing gas exchange surface. (K) Schematic of experimental design for LPS-induced lung injury. Rhesus macaque dams received intra-amniotic LPS injection at GD130, with subsequent fetal delivery by C-section at 16h (n=8), 48h (n=3), or 5d (n=7) after injury. (L-Q) Extensive disruption of gas exchange surface, shown through the disordered patterning and interactions of HOPX^+^ AT1 cells (red) and EDNRB^+^ aerocytes (green) in both mild and severely injured regions up to 5d following LPS. R) Model of alveolar simplification following LPS injury. Control Rhesus fetal lung exhibits organized, complex patterning of epithelial (AT1, AT2), mesenchymal (myofibroblasts [MyoFB], matrix fibroblasts [MatrixFB]) and endothelial (capillary endothelium [Cap endo]) cells with formation of new alveolar septae. This patterning is disrupted in LPS-injured lungs, with increased myeloid cell and neutrophil infiltration, and subsequent mesenchymal response (evidenced through increased inflammatory fibroblasts (InflamFB) ultimately leading alveolar simplification and loss of alveolar septal formation. (Scale bars = 50 μm). For full list of samples and treatment groups, see Supplemental Table 1. *(ASMA = α-Smooth Muscle Actin [myofibroblast marker]; EDNRB = Endothelin Receptor B [alveolar capillary/aerocyte marker]; HOPX = Homeobox Only Protein [AT1 marker]).*

Trimesters in Rhesus are approximately 55 days, compared to 90 days in humans; therefore, GD105 equates to approximately 25-26 weeks of human gestation, GD130 to approximately 30-32 weeks, and GD150 to 35-36 weeks^34^ (Figure 1A). Morphologically, GD105 represents the transition from late canalicular to early saccular lung, while GD130 represents late saccular to early alveolar stage, and extensive alveolarization is evident by GD150 (Figure 1B-J, S2C,I,O). During alveolarization, alveolar septal structures (AS) expand as the gas exchange surface develops; epithelial and endothelial lineages form close interactions along the extending AS, preparing the lung for birth and air breathing (Figure 1H-J). As alveologenesis proceeds, AT1 and AT2 cells (Figure 1E-G) are progressively specified while alveolar structural complexity increases with septal maturation. Recent studies in mouse and human have defined EDNRB^+^ endothelial cells as specialized alveolar capillaries (also called aerocytes) which are critical for gas exchange^35–37^. Similarly, progressive maturation of the alveolar capillary network (Figure 1H-J) with increasingly close association and co-localization with AT1 cells drives development of the primate gas exchange surface. Histological analysis (Figure S2J-P) and bulk RNA sequencing (Figure S3A-H) demonstrate high concordance of proliferative and TF expression dynamics with murine models of lung development, but with structural and temporal similarities to human lung development (Figure S1A-C). Therefore, Rhesus macaque is a translationally relevant model of lung development, with GD130 representing a period of active morphological and cellular maturation as alveolar septae and capillary networks form.

Most 26-week and later premature human infants can be successfully resuscitated after birth and supported in the neonatal intensive care unit, so human tissue obtained at these developmental times comes from small numbers of infants with severe non-pulmonary disease or after medical intervention^38^. Given these factors, Rhesus provides a unique model to perform controlled experiments focused on understanding mechanisms of primate developmental lung injury. In order to model fetal chorioamnionitis, we treated Rhesus fetuses at GD130 with intra-amniotic injection of endotoxin/lipopolysaccharide (LPS), an injury that has been shown to mimic the inflammation present during chorioamnionitis in prenatal mouse, rabbit, pig, and sheep^32,33,39,40^, as well as Rhesus macaque in prior studies from our group^33,39,40^. LPS robustly induced IL1β, TNFα, and IL-6 after 16h in both lung tissue and alveolar wash in Rhesus^39,41^ and led to rapid infiltration and activation of neutrophils and myeloid cells in the fetal Rhesus lung^41^. A detailed evaluation of the inflammatory state of the chorioamnionitis lung in Rhesus macaques is reported seperately^41^. Here, we evaluated the hypothesis that this inflammatory milieu would directly impair the developmental progression of the Rhesus lung.

By 16h after LPS treatment (Figure 1K), significant lung injury is evident (Figure 1L-M, S4A, S5A-B, G-H) with areas of both mild and severe disruption of tissue architecture, recapitulating the heterogeneous lung damage seen in human infants after chorioamnionitis. Inflammation impairs the close opposition of AT1 cells and aerocytes in both mildly and severely injured regions of the lung (Figure 1L-Q, S5G-L) and causes loss of alveolar septal structures most obvious at 48h after injury (Figure 1P-Q, S4B, S5). More alveolar septae are detectable at 5d post-LPS (Figure 1P-Q, S4C, S5 E-F, K-L), primarily in more mildly injured regions (Figure S5E,K). In more severely injured regions, we also observed loss of differentiated epithelial cells (Figure S5A-F). By 5d after LPS injury, differentiated AT1 and AT2 markers were only partially restored (Figure S5E-F). Overall, in both mildly and severely injured regions, there is significant alveolar simplification secondary to inflammatory injury (Figure S4A-C). Since chorioamnionitis and prematurity are known to increase risk of secondary viral infection^29^, we also assessed the expression of the SARS-COV2 priming protease TMPRSS2 in control and injured lungs (Figure S6), and found no significant expression, even with inflammatory stress, concordant with recent mouse and human data^42^. Together, our findings suggest that LPS-mediated inflammatory injury causes significant, multifocal, and persistent injury to the developing primate lung, with major shifts in cell quorum and tissue architecture (Figure 1R), resulting in pathological findings similar to those seen in premature infants after lung injury^24^.

### A single cell atlas of developmental lung injury

To further evaluate pulmonary tissue remodeling and develop insight into the pathophysiology of the perinatal lung injury, we performed single cell RNA sequencing (scRNA-seq) on GD130 fetal lungs from control animals and 16h after LPS treatment (Figure 2A). Then, to specifically evaluate injury responses in the developing gas exchange surface of the primate lung, we performed a focused analysis of epithelial, endothelial, and mesenchymal cell populations (Figure 2B-C).

**Figure 2.**
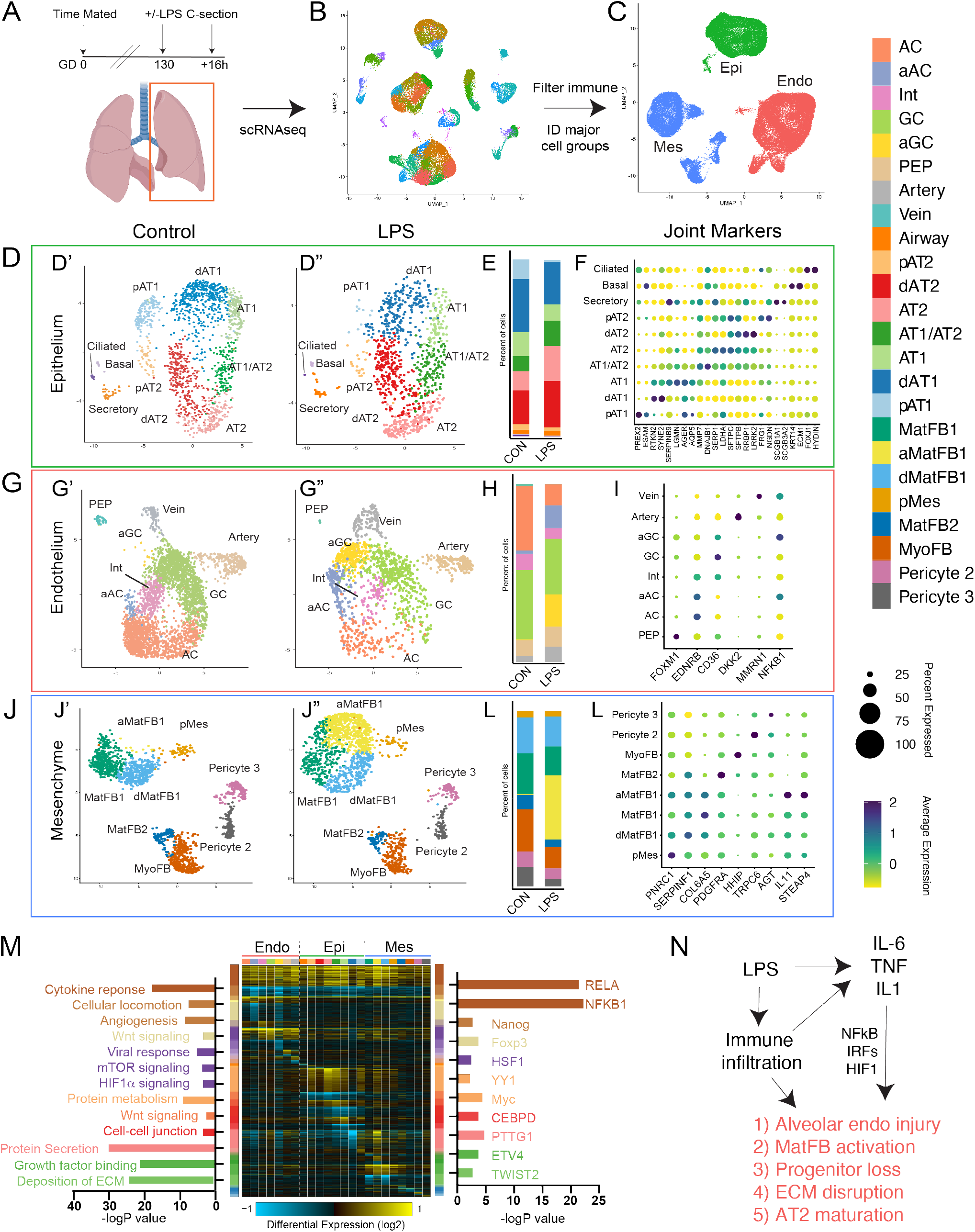
Single cell interrogation of developmental lung injury. A-C) scRNAseq of control (n=2) and LPS-treated (n=2) primate lungs. UMAP projection of all cells was generated in Seurat (B), and major structural cell populations were identified by expression of PECAM1/CD31 (Endo), CDH1 (Epi), or COL1A1 (Mes) (C) for additional analysis. D-F) UMAP projection of epithelial cell clusters from control (D’) and LPS-treated (D”) Rhesus. Colors are coded as per the sidebar, except that Airway is separated into ciliated, basal, and secretory cells. Relative proportions of each cell type are shown in (E). Relative frequency of cells expressing cell markers in (F), identified based on cellular enrichment and comparison to marker gene expression in LungMAP datasets^21,25^. G-L) Endothelial cell (G-I) and mesenchymal cell (J-L) UMAP projection, frequency, and cell markers. M) cellHarmony heatmap of injury vs. control differentially expressed genes (rows) for these cell populations (columns), highlighting GO, Pathway, and experimentally observed TF-target gene-module enrichment. Cell types are color coded as indicated in the sidebar. Colors of bars are based on differentially expressed gene sets on Y axis. N) Model of overall LPS effect combining data from Figures 1&2. *AC = alveolar capillary, aAC = activated AC, Int = intermediate capillary, GC = general capillary, aGC = activated GC, PEP = proliferative endothelial progenitor, MatFB = matrix fibroblast, aMatFB = activated matrix fibroblast, dMatFB = differentiating matrix fibroblast, MyoFB = myofibroblast, pMes = proliferative mesenchyme, AT2 = alveolar type 2, pAT2 = progenitor AT2, dAT2 = differentiating AT2, AT1 = alveolar type 1, pAT1= progenitor AT1, dAT1 = differentiating AT1, ECM = extracellular matrix, CON = control*.

Clusters for subtypes of epithelium (Figure 2D-F), endothelium (Figure 2G-I), and mesenchyme (Figure 2J-L) were identified using Seurat v3^43^, with parameters set to identify major cell populations rather than multiple similar subclusters. We used LungMAP^38,44^ and Human Cell Atlas^45^ annotations from human lung to determine cell type identity on the basis of enriched expression of key marker genes. We detected the major cell types described in human studies in all three populations, with highest similarity to early life human datasets^38,44^.

In the epithelium (Figure 2D-F), we detect the major cell types of both the airway and alveolar regions. In the alveolus, there are multiple distinct cellular states of AT1 and AT2 cells, and a cell population expressing markers of both cell types similar to the AT1/AT2 cell state described during murine lung development^46^ and in a recent human cohort^38^. Recent lineage tracing results have suggested that AT1 and AT2 progenitor populations diverge early in mouse lung development, with distinct AT1- and AT2-defined clones emerging as early as E13.5 and becoming increasingly common as development proceeds; single cell RNA sequencing enabled separation of AT1 and AT2 progenitors from more fully differentiated distal epithelium by E17.5^47^. Similarly, there were distinct, separable progenitor AT1 and AT2 lineage clusters in prenatal Rhesus. Pseudotemporal lineage inference (Figure 3A)^48^ demonstrated differential trajectories supporting AT1 and AT2 cell evolution from these progenitors to fully differentiated cells through distinct differentiation intermediates. Lineage inference suggested that the AT1/AT2 cell state derives preferentially from the AT2 lineage branch, and predicts some AT1 differentiation through this state, in accordance with multiple studies in mice that AT1 cells can arise from AT2 progenitors^49–51^.

**Figure 3.**
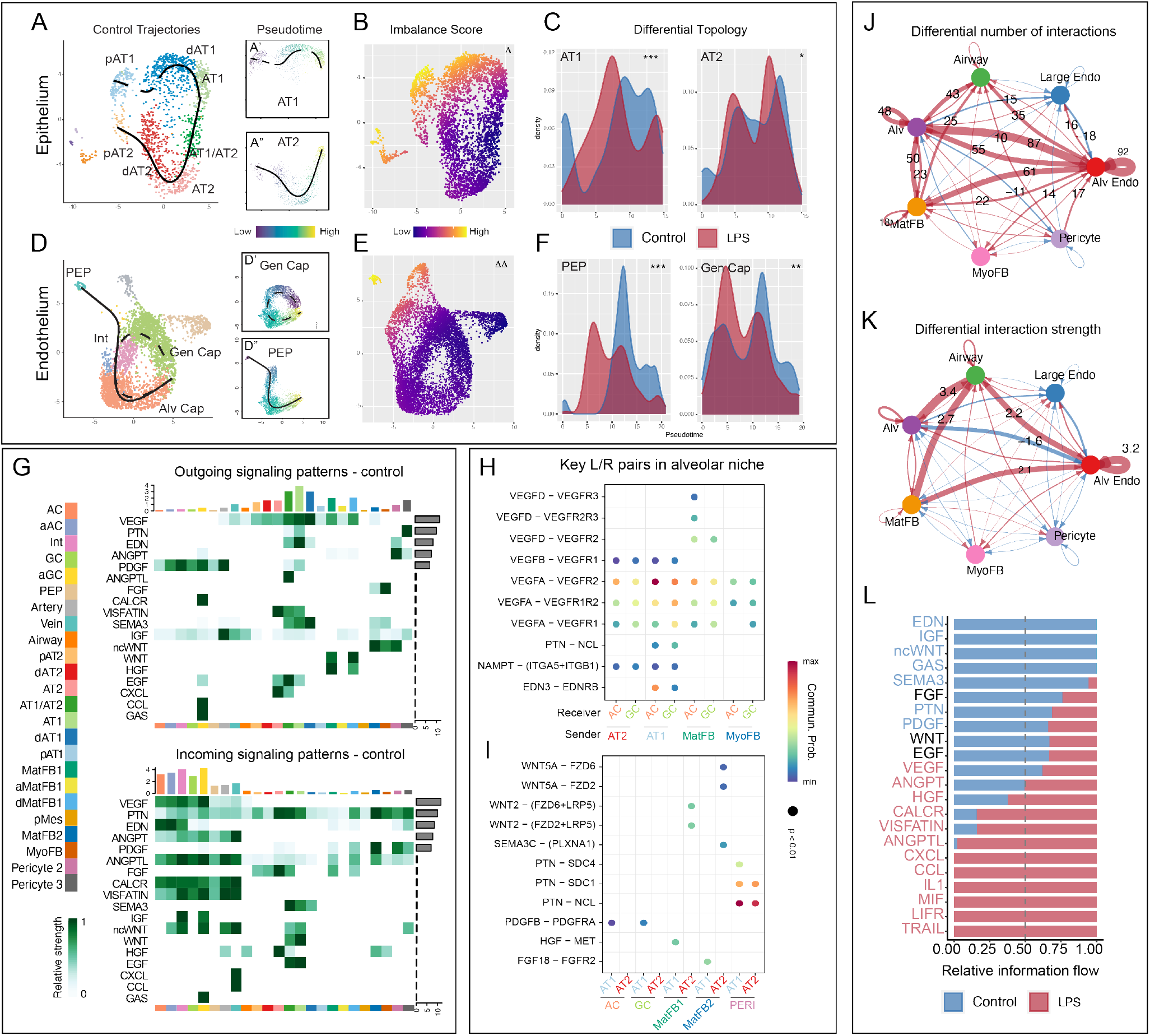
Inflammatory injury disrupts progenitor cell differentiation and niche signaling in the developing alveolus. Differential trajectory analysis of differentiation of alveolar epithelial (A-C) and endothelial (D-F) lineages with LPS injury. Control epithelial trajectories (A) obtained with *Slingshot* show AT1 and AT2 trajectories arising from separable progenitor lineages. Alveolar capillaries (Alv Cap) arise from both PEPs and general capillaries in control (D). LPS causes significant transcriptional imbalance in both epithelium (B) and endothelium (E) with disruption of developing AT1 (C) and alveolar capillary trajectories (F). Differential trajectories were calculated using *Slingshot* and *Condiments*, Δ = p <1e-10 and Δ Δ = p <1e-13 for differential trajectory topology. *=p<1e-3, **=p<1e-10, ***= p<2.2e-16 for differential trajectory progression. G-L) CellChat analysis of signaling milieu of the developing alveolus. G) Major ligand/receptor (L/R) relationships in the developing primate alveolus displayed as a heatmap showing outgoing ligands (G top panel) and incoming receptivity (G bottom panel). Overrepresented signaling pathways are along left Y axis, and cell identity is along bottom X axis, with colors corresponding to cell identity as per the sidebar. Right Y axis shows relative strength of the predicted signaling pathway in the overall network, and top X axis shows relative contribution of the cell population to the overall signaling milieu. H-I) Key ligand and receptor interactions targeting aerocytes and general capillaries (H) or AT1 and AT2 cells (I). Ligand-receptor pair is shown along Y axis, with sender and receiver cells along X axis. J-L) Analysis of changes in signaling milieu of the LPS-treated lung. Blue indicates increased signaling in control, while red indicates increased signaling with LPS. Significant increases in the number (J) and strength (K) of interactions occurs with LPS, with shift (L) from developmental signaling pathways enriched in the baseline alveolar niche to increased inflammatory signaling in the damaged alveolar niche. Blue text in (L) indicates enriched signals in control, and red indicates enriched signals in LPS. *Cell identity abbreviations are the same as in Figure 2, and standard gene names and abbreviations are used for signaling pathways, ligands, and receptors.*

In addition to arterial and venous endothelial cells, we identified three alveolar capillary endothelial lineages, which correspond to alveolar and general capillary populations described during mouse and human development and disease^35–37^ as well as an intermediate population expressing markers for both aerocytes and general capillaries (Figure 2C). We also identified a separate endothelial population, characterized by high level proliferation (despite normalization for cell cycle genes) and a distinct progenitor gene signature (Figure 2G-I, 3D, S7). Pseudotime analysis predicts that this population functions as a progenitor lineage for both alveolar and general capillaries (Figure 3D). Comparison of the gene signature and pseudotime trajectory of these cells to previously reported single cell data sets from mouse development^52,53^ and early human development^54^ suggests that this population, marked in our studies by FOXM1, is a conserved proliferative endothelial progenitor (PEP) lineage for the lung (Figure S7A-K). We observed mesenchymal heterogeneity (Figure 2J-L) similar to that reported in both mouse and human, with multiple populations of pericytes, matrix fibroblasts, and myofibroblasts, though we do not detect a distinct lineage of smooth muscle cells, possibly owing to relatively lower representation of proximal airway in our dataset.

Global analysis of differential gene expression of the injured versus control lungs (Figure 2M) demonstrated extensive activation of inflammatory pathways in all cellular populations, with generalized activation of NFKB1 and RELA targets consistent with active NFκB signaling throughout the structural cells of the lung. Detected gene expression differences were common among related cell types, and are consistent with our previous observation from mouse lung development that NFκB is an important signaling mechanism regulating the timing of lung maturation^55^. We also noted loss of progenitor lineages in both the epithelium and endothelium (Figure 2E,H), with reduction in quorum of AT1 and AT2 progenitors as well as PEPs in injured lungs. AT2 cell maturity markers increased, especially secretory and ER-related GO terms in remaining AT2 cells (Figure 2D-F,M), consistent with prior reports indicating that inflammation promotes AT2 maturation^40,56^. Aerocytes and general capillaries showed significant inflammatory activation with a shift of transcriptional prolife in both cell populations to an activated state (Figure 2G-I), characterized by expression of HIF1α-mediated hypoxia target genes and loss of angiogenic factors (Figure 2M). Matrix fibroblasts, particularly type 1 matrix fibroblasts, also showed significant response to LPS, taking on an inflammatory state marked by high level expression of IL1 and TNF target genes (Figure 2D-F,M). These findings demonstrate that intra-amniotic LPS causes extensive damage to the developing gas exchange surface of the primate lung (Figure 2N).

### Disrupted patterning of the primate gas exchange surface during prenatal injury

Next, we evaluated the impact of LPS injury on developmental trajectories of key progenitor populations in the epithelium and endothelium during alveolar patterning. Pseudotime inference can predict developmental trajectories, but one limitation of this methodology has been comparison of differential trajectories between experimental perturbations; the recently reported package *Condiments*^57^ addresses this deficiency by directly evaluating changes to trajectories and along specified trajectories between experimental conditions. We used this tool, in combination with *Slingshot* pseudotime inference^48^, to directly evaluate the impact of LPS injury. In the epithelium, the AT1 trajectory is significantly disrupted by LPS, with loss of AT1 progenitors and reduction in fully differentiated AT1 cells (Figure 2D, 3A-C). Conversely, the AT2 trajectory is only mildly disrupted, with loss of AT2 progenitors but compensatory distal differentiation along the wild-type trajectory (Figure 2D, 3A-C), consistent with extensive evidence that perinatal inflammation promotes AT2 maturity^40,56^. In the endothelium, differentiation of aerocytes is reduced, induced by loss of FOXM1^+^ PEPs (Figure S5G-L, S7G-I) and decreased differentiation from general capillaries, with preferential accumulation of EDNRB^+^/APLNR^+^ intermediates in both trajectories (Figure 3D-F). There is also a clear transition of both aerocytes and general capillaries to an activated, IL1- and TNF-responsive state (Figure 2G-I). Therefore, concordant with histological data, trajectory analysis confirms that a predominant effect of LPS on the developing lung is disorganization of critical AT1-alveolar capillary interactions.

We then performed global signaling analysis of control and LPS-injured lung using the package CellChat^58^. CellChat uses cell-specific expression of ligands and receptors to generate a network comprising the cell signaling milieu of a system, and then identifies key overrepresented pathways within that system. We applied this method to decipher the signaling niche of the developing alveolus and define how these communication patterns were disrupted by inflammatory injury (Figure 3G-L). In general, major signaling pathways known to pattern distal lung development in mice are recapitulated in Rhesus, with cellular expression domains and receptivity that match previous data from other systems. For example, the WNT signaling pathway is defined by expression of WNT2 ligand from a subset of matrix fibroblasts, while WNT5A is expressed primarily in myofibroblasts and pericytes; signaling response to both signals is prominent in alveolar epithelial and endothelial lineages, as expected from data in mice and human precision-cut lung slices^49,59,60^. CellChat estimates the contribution of individual pathways to the overall signaling milieu; we identified vascular endothelial growth factor (VEGF), pleiotrophin (PTN), endothelin (EDN), and platelet-derived growth factor (PDGF) signaling as major contributors to alveolar maturation (Figure 3G). Closer evaluation of these interactions shows that VEGFA and EDN3 ligand from developing AT1 cells signal to developing alveolar capillaries, likely contributing to the patterning of the alveolar gas exchange surface (Figure 3H). Contemporaneously, WNT, PTN, FGF, and HGF signals from mesenchymal cells promote epithelial maturation (Figure 3I). Matrix fibroblasts function as a major signaling hub in the mesenchyme, while myofibroblasts, which appear to be mechanically important in dynamic maturation of alveolae^61^ (Figure 1B-D) are less prominent in the enriched signaling network. Together, these data define an alveolar signaling niche, containing both conserved signaling interactions (e.g. WNT, FGF) and novel interactions (e.g. EDN), driving the patterning of the alveolar gas exchange surface in primates.

This alveolar signaling niche was significantly disrupted after inflammatory injury by LPS. Differential interaction analysis demonstrated that both the number and strength of interactions within the lung increased after LPS injury, with significantly more interactions between alveolar epithelium, endothelium, and mesenchymal lineages (Figure 3J-K). While growth factors and maturation signals dominate the control signaling milieu, LPS induces significant increases in CCL, CXCL, IL1, and TNF signaling pathways, consistent with a generalized inflammatory injury to the lung (Figure 3L). Corresponding loss of major developmental signals including EDN, IGF, PTN, and WNT implied significant disruption of the developing AT1-aerocyte network (Figure 3H-L). These findings are concordant with the observed histological disruption of the gas exchange surface and emphasize that developing aerocytes are a major target of injury during late gestation. In the mesenchyme, we observed that the large, new population of activated matrix fibroblasts (Figure 2J) appeared to elaborate pro-inflammatory signaling in the injured lung (Figure S8A-B), with robust expression of CCL and CXCL chemokines and decreased expression of WNT and HGF ligands. The net effect of these changes is a signaling milieu dominated by inflammatory ligands and with reductions in major developmental pathways. These findings imply that inflammatory injury, both to the signaling niche of the lung and the developmental trajectory of key progenitors, is a major mechanism causing lung injury in prenatal Rhesus, driving the alveolar simplification evident at 5d post LPS.

### Combined blockade of IL1 and TNF signaling prevents inflammatory lung injury

Therefore, we proceeded to test the hypothesis that direct blockade of inflammatory pathways driving experimental chorioamnionitis could protect the developing lung from injury. We treated pregnant Rhesus dams with anakinra (100mg, Kineret, SOBI), a potent IL1 receptor antagonist, and adalimumab (40mg, Humira, AbbVie Inc), an anti-TNF monoclonal antibody, by subcutaneous injection 3h prior to LPS exposure, and by intra-amniotic injection (anakinra 50mg and adalimumab 40mg) 1h prior to LPS treatment (Figure 4A). This combination blockade did not reduce either myeloid or neutrophil accumulation in the lung compared to LPS, but modestly reduced inflammatory cytokine levels in the BAL of treated animals and the expression of TLR, IL1, and TNF response pathways in several immune lineages (finding detailed in^41^). Despite these modest changes, inflammatory blockade provided remarkable protection to lung alveoli (Figure 4B-S). Cellular quorum by scRNAseq returned to near baseline levels, with both epithelial (Figure 4B-C) and endothelial progenitors (Figure 4D-E) observed at control levels. Concordant with this observation, anti-inflammatory blockade significantly improved lung injury severity, with many regions nearly indistinguishable from uninjured lung (Figure 4H-S). Histological quantification demonstrated normalization of septal numbers and injury score to control levels (Figure 4T-U, Figure S9A-C). Additionally, nearly 80% of differentially regulated genes in LPS lungs were normalized after blockade (Figure S9D-E). CellChat analysis demonstrated significant improvement in major signaling pathways including EDN, PTN, FGF, and HGF after TNF and IL1 blockade (Figure 4F-G). Despite these changes, a distinct inflammatory signal was still present in treated lungs compared to control, suggesting that the inflammatory blockade provided by anakinra and adalimumab was partial (Figure 4G).

**Figure 4.**
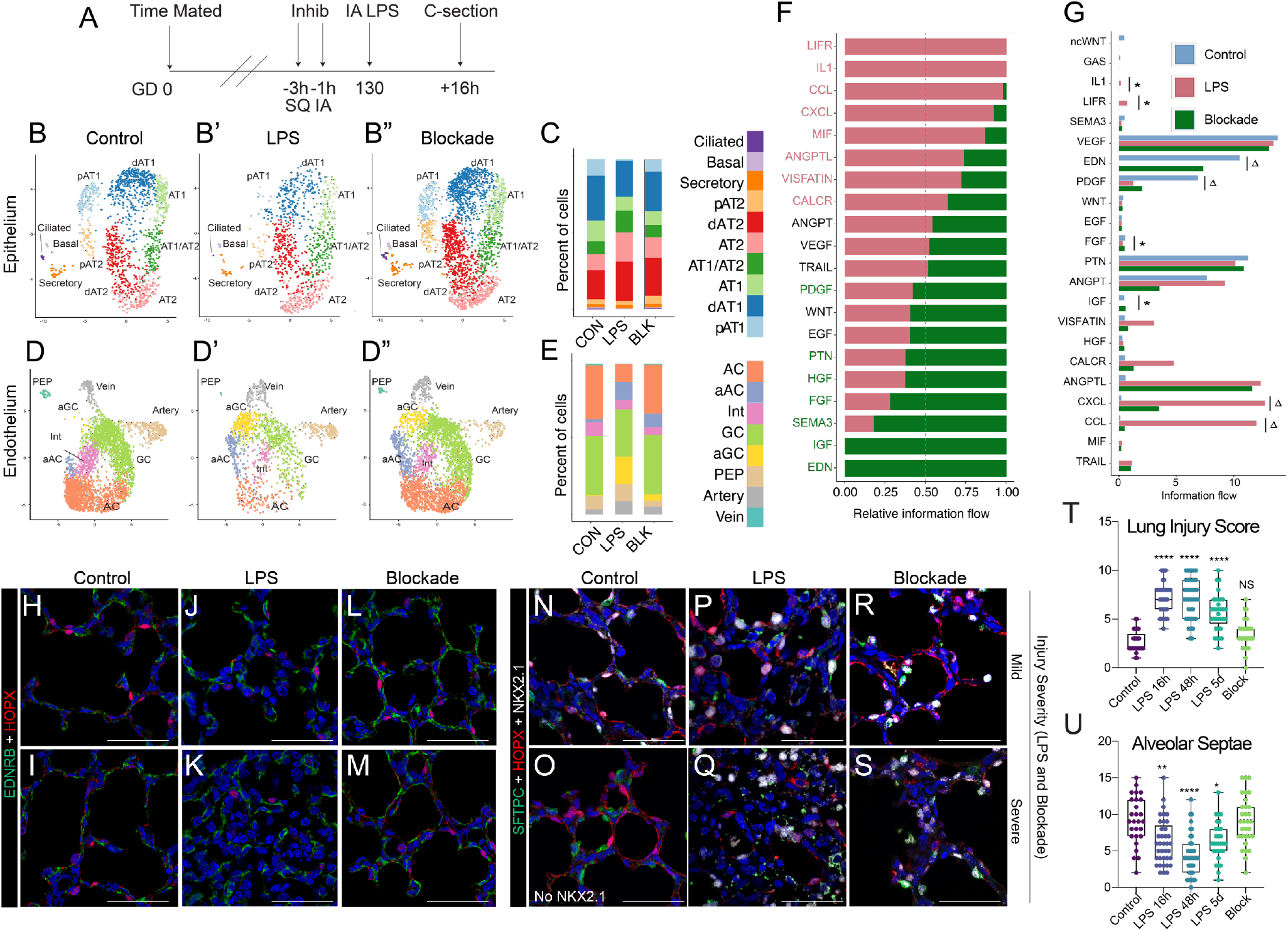
Combined IL1 and TNF blockade protects the developing lung from inflammatory injury. A) Schematic of experimental design. Rhesus dams received adalimumab and anakinra subcutaneously (SQ) 3h prior to LPS and by intra-amniotic injection (IA) 1h prior to LPS. They then received IA LPS and were examined at 16h post treatment (n=5; n=3 for scRNA-seq). Data for LPS and control animals is shown for comparison. B-E) UMAP projection of epithelial (B) and endothelial (C) cell clusters from control (B, D), LPS-treated (B’, D’) and combined blockade (B”, D”) animals. Colors are coded as per the sidebar. Relative proportions of each cell type are shown in (C) and (E). Combined blockade prevents loss of epithelial and endothelial progenitor lineages and blunts activation of inflammatory pathways in alveolar capillary endothelium. Cell proportions in both epithelium and endothelium resemble control animals. F) CellChat analysis of signaling milieu of the alveolus after combination blockade compared to LPS. Red text in indicates enriched signals in LPS, and green enriched signals in combined blockade. The lung stromal signaling milieu reverts markedly compared to LPS treatment. Notably, FGF, HFG, pleiotrophin, and endothelin signaling improve with inflammatory blockade. G) Comparison of intensity of major signaling pathways in control, LPS, and blockade conditions demonstrates some pathways normalize to baseline (not significantly different than control, denoted with *) while others improve but do not normalize (significantly improved from LPS, but significantly elevated compared to control, denoted with ^Δ^), including most inflammatory pathways. H-M) Combination blockade prevents disruption of AT1/alveolar capillary interactions. N-S) Maintenance of differentiated epithelial cells in alveoli increases after combination blockade. T-U) Quantifications of lung injury score^60^ and septal numbers show normalization compared to control and significant improvement compared to LPS. * = p <0.05, ** = p <0.01, **** = p < 0.0001 by ANOVA compared to control. *Abbreviations as above.*

These results demonstrate that the combination of IL1 and TNF blockade is sufficient to prevent severe lung injury in experimental chorioamnionitis, despite continued cellular recruitment of immune cells and significant levels of inflammatory cytokines in the BAL^41^. We therefore hypothesized that blockade may reduce inflammatory activation of resident stromal and immune cells in the lung, preventing secondary injury to lung structure. To identify immune-stromal signaling relationships in LPS and treatment conditions, we used our unfiltered scRNAseq data set, containing both structural and immune cell lineages, and integrated cells from all captures to identify the predominant populations (Figure 5A). We then performed CellChat analysis, nothing that myeloid immune lineages, including monocytes, macrophages, and dendritic cells, formed a major signaling partner with structural cells in both LPS and blockade conditions (Figure 5B-C).

**Figure 5.**
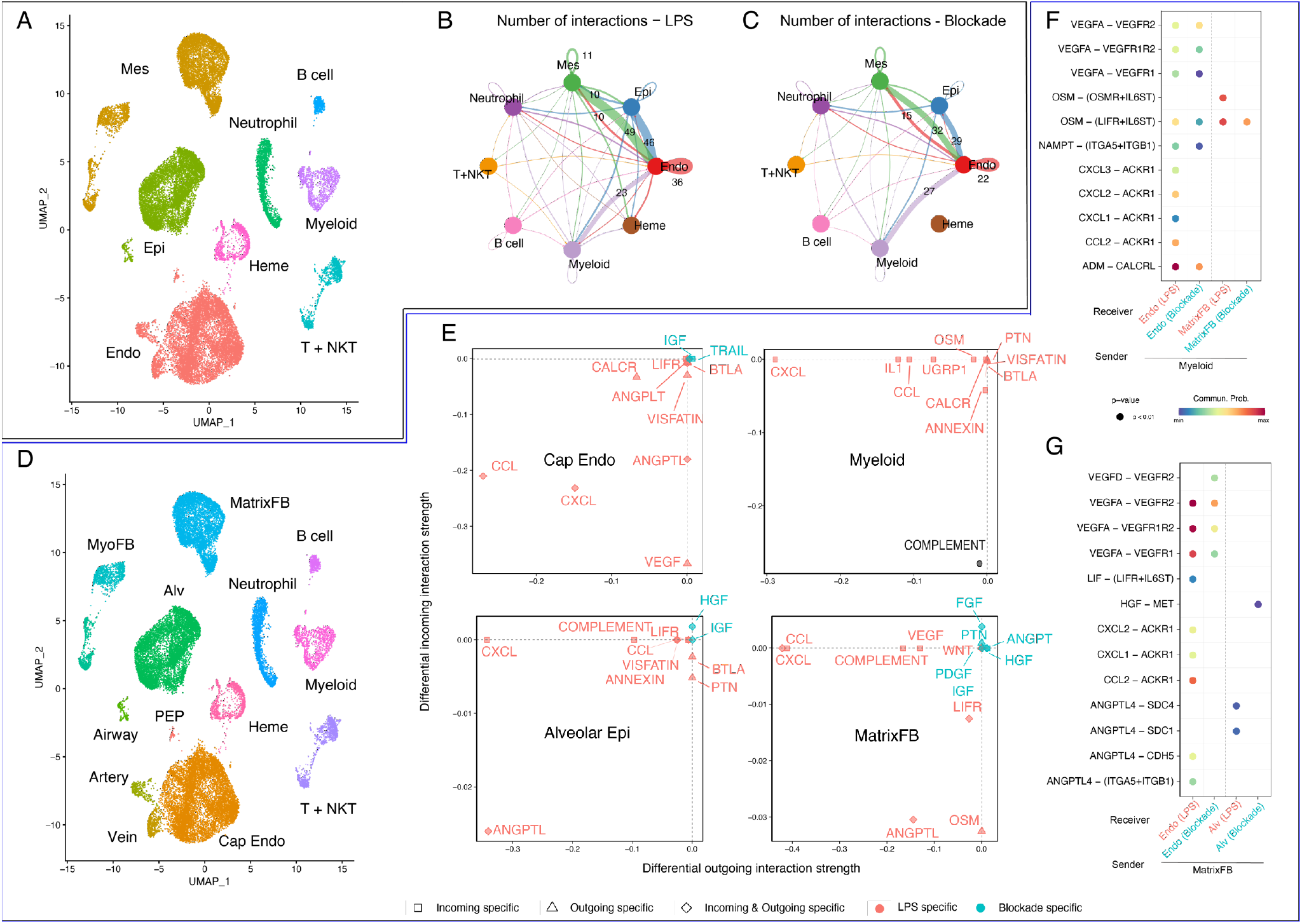
Combined IL1 and TNF blockade prevents inflammatory injury to the alveolar signaling niche via modulation of CCL and CXCL cytokines in matrix fibroblasts and innate myeloid cells. A) UMAP projection of large cell populations including immune cells used to identify major interactors in LPS vs blockade lungs. CellChat analysis demonstrates that myeloid immune lineages signal extensively to epithelium, endothelium, and mesenchyme in both LPS (B) and blockade (C) conditions. D) UMAP projection of refined cell populations used for analysis in (E-G) E) CellChat differential response analysis show that myeloid cells and matrix fibroblasts elaborate extensive CXCL and CCL ligands in response to LPS (red), with significant reduction in this inflammatory activation after combination blockade (blue). Notably, blockade increases expression of key developmental niche signals from the matrix fibroblasts (blue). F-G) Major differential ligand and receptor pairs involved in myeloid signaling to endothelial and mesenchymal cells (F) and activated matrix fibroblast signaling to endothelial and alveolar endothelial cells, showing intensity with either LPS (red) or blockade (blue.) *Mes = mesenchyme, Epi = epithelium, Endo = endothelium, Heme = hematopoietic, T + NKT = T and NKT cells, MyoFB = myofibroblast, MatrixFB = matrix fibroblasts, PEP = proliferative endothelial progenitor, Alv = alveolar epithelium, Cap endo = capillary endothelium (inclusive of general capillaries and aerocytes).*

Next, we subdivided structural cells into major subcategories (Figure 5D) and found that myeloid cells and activated matrix fibroblasts produced extensive inflammatory signaling modulators, including CXCL and CCL ligands, following LPS treatment (Figure 5E-G). CXCL and CCL pathway response is predicted in alveolar endothelial and epithelial cells via atypical chemokine receptor 1 (ACKR1)^62^ after LPS, likely contributing to the significant disruption of epithelial and endothelial patterning seen in these animals. ACKR receptors are important modulators of cellular inflammatory response, where they both promote endothelial adhesion of immune cells^63^, typically in venules, and regulate cell proliferation and differentiation^64^. Global knockout of ACKR1 in mice dramatically reduces lung inflammation following systemic LPS challenge^65^. Adrenomedullin (ADM) signaling from myeloid cells to alveolar endothelium via CALCRL signaling also after LPS; recent mouse data has implicated ADM-CALCRL signaling as an important factor in repair of the neonatal lung after hyperoxia injury^66,67^. Blockade of IL1 and TNF signaling dramatically blunts CCL and CXCL cytokine expression in both matrix fibroblasts and myeloid cells (Figure 5F-G), and response to these ligands correspondingly decreases throughout alveolar cells. ADM-CALCRL signaling decreases to near control levels in blockade (Figure 4G). Remarkably, matrix fibroblasts elaborate FGF, HGF, and IGF ligands (Figure 5E,G) preferentially after inflammatory blockade, similar to the milieu of the control alveolar niche, restoring a developmental pattern of response to these signals in the epithelium and endothelium.

Taken together, these data support a model of developmental lung injury (Figure 6A-B) in fetal primates whereby LPS-induced inflammatory cytokine signaling directly injures the structural and signaling niches of the developing alveolus. This altered signaling milieu then causes widespread activation of inflammatory pathways, promoting a pro-inflammatory state of myeloid immune lineages resident in or recruited to the lung. Together, these inflammatory signals potentiate injury to the alveolar niche through elaboration of CCL and CXCL ligands, ultimately leading to failed alveologenesis and alveolar simplification. Blockade of IL1 and TNF signaling is sufficient to blunt this pro-inflammatory state, decreasing cell-intrinsic inflammatory response in structural cells and promoting a more tolerogenic phenotype in myeloid cells, leading to protection of the developing alveolus, maintenance of key developmental signaling relationships, and functional development of the gas exchange surface in preparation for birth and post-natal respiration (Figure 6C).

**Figure 6.**
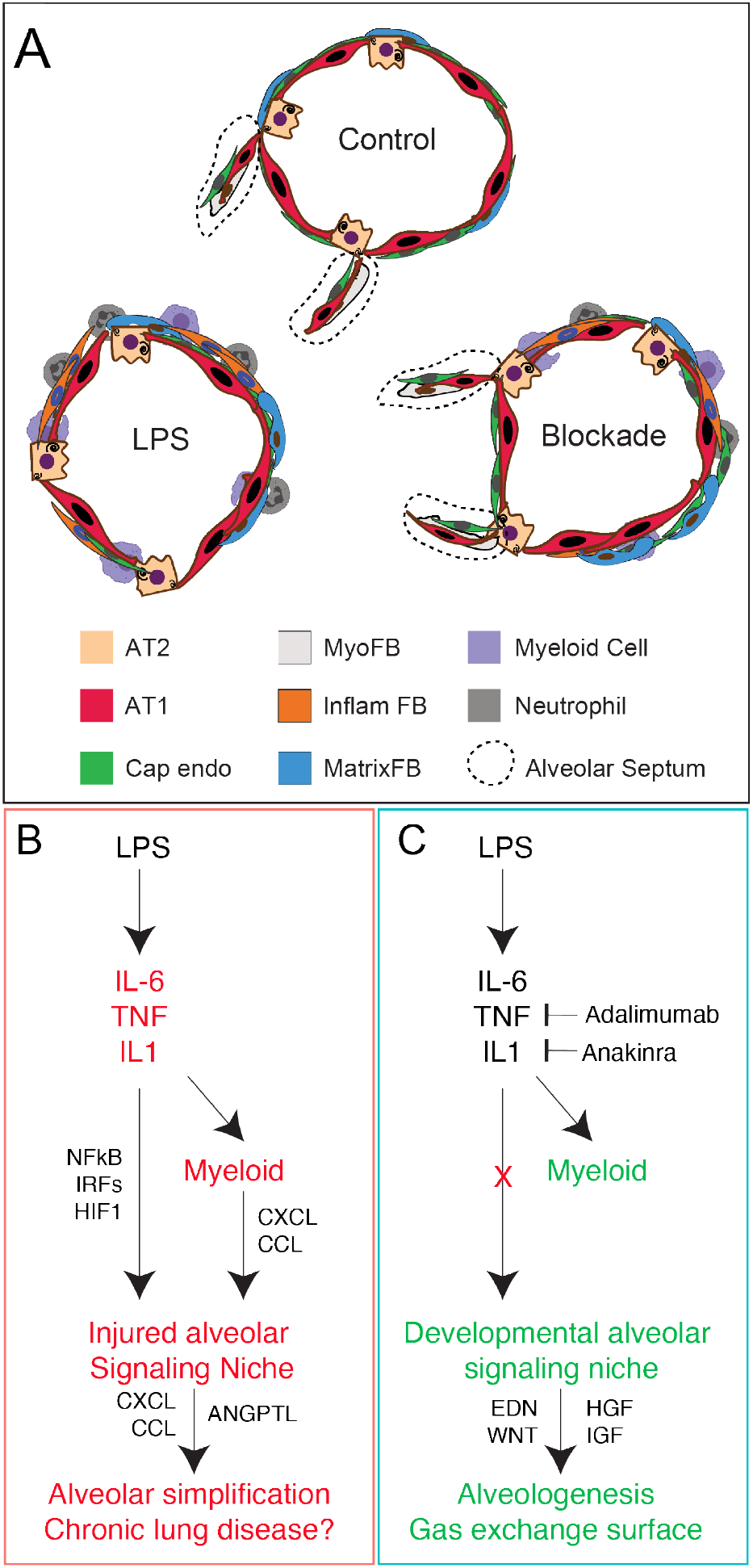
Model of pathogenesis and treatment of inflammatory lung injury in macaque. A-B) LPS causes significant alveolar simplification via activation of global inflammatory cytokines including IL1, IL6, and TNF, leading to activation of these pathways across the alveolar niche. Myeloid cells potentiate this injury via expression of CXCL and CCL cytokines, which are also elaborated from injured matrix fibroblasts, worsening injury. Inflammatory blockade blunts this pro-inflammatory state, reduces CCL and CXCL cytokine expression, and protects the signaling milieu of the alveolus dominated by developmental growth factors and morphogens, thereby promoting continued alveologenesis. *Abbreviations as previously used.*

## Discussion

### A conserved niche for mammalian alveologenesis

Successful adaption to post-natal life is entirely dependent upon the formation and function of the pulmonary alveoli which mediate the efficient exchange of oxygen and carbon dioxide. Much of what we know about alveologenesis comes from decades of controlled mouse experimentation, which has generated detailed knowledge regarding signaling interactions and transcriptional regulators active during lung morphogenesis^4^. Nonetheless, key differences between mouse and human lung limit what can be learned from purely murine studies. Nonhuman primates can be used for controlled experimentation, as in the current study, to directly assess causality; such causal studies are rarely possible in humans. Here, we utilized the Rhesus model to define the alveolar niche in primates. Rhesus shares key morphological and developmental timing with human lung development, and our data demonstrate conservation in key signaling factors and cell signatures. Major markers of alveolar epithelial cells^46,47,49^, mesenchymal lineages^68–71^, and endothelial specialization^35,36,53^ are similar to what has been defined during mouse development and in human lung datasets^17,38,44,45^. This high-level conservation between mouse, primate, and human lung emphasizes the requirement for functional alveologenesis in mammalian physiology. With strong connections to both mouse and human physiology and expression, the Rhesus macaque model provides a unique opportunity to bridge murine and human studies. While recent data has suggested some drift in adult lung expression patterns and signaling between mouse and human^17^, our study suggests important evolutionary conservation during alveologenesis and may provide a framework for understanding perturbations in alveolar development beyond inflammatory injury. Given that endothelial-AT1 interactions are central to the development of the alveolus, and are directly injured by LPS, identifying determinants of this co-development is a high priority that may allow development of new therapeutics for both pediatric and adult lung diseases.

Beyond cellular and structure conservation, our data also defines a dedicated niche of signaling factors and gene networks which control the growth and patterning of the developing primate alveolus. WNT^49,59,60^, FGF^72–74^, EGF^75^, and VEGF^35^ signaling, all of which are critical in mouse alveolar development, function in multiple signaling loops in the Rhesus lung during normal development and are heavily disrupted during injury, implying important functional conservation. Our data also provides new insight into alveolar signaling, as we also observe centrality of several less studied pathways in the niche. EDN and PTN signaling in AT1-aerocyte interactions are particularly prominent pathways which merit further study in Rhesus and other systems. Adaptation of techniques successfully used in human and mouse development, including organoids^15,50^ and precision-cut lung slices^59,76^ will be important in future studies to provide additional specificity in our understanding of these pathways, and identification of key signaling interactions that differ between mouse and primate.

### Immune signals as important regulators of lung development and repair

A provocative aspect of our study is the finding that inflammatory blockade is sufficient to protect the developing lung from significant injury following LPS treatment. These data add to an evolving understanding of the role of immune cells in both lung development and lung repair. Recent data in adult mice has demonstrated that inflammatory cytokine pathways, particularly IL1^77,78^, functionally promote lung regeneration after multiple types of injury, driving AT2 proliferation^77^ and AT1 differentiation through a regenerative transition state^78^. Myeloid lineages have been reported to promote regeneration after pneumonectomy via CCR2/CCL2 signaling^79^. We observe the opposite effect in the developing primate lung, where IL1 and TNF blockade protects the lung from injury, and CXCL/CCL chemokine signaling is a major hallmark of the LPS-treated Rhesus lung that is suppressed by IL1 and TNF blockade.

Our data suggest several possibilities to explain these differences. First, they could relate to tissue and immune maturity. Recent reports demonstrate persuasively that the perinatal immune system differs significantly from adult immunity^80,81^. Fetal immunity is carefully balanced to provide protection from pathogens while also allowing development of tolerance and colonization of microbiota. Extensive data from human studies, and our companion report on Rhesus immune development^41^, shows that chorioamnionitis disrupts this immune development^32,33^. Clinical chorioamnionitis is typically treated with antimicrobials, which also have unintended but significant impacts on immune development^82,83^. Each of these individual impacts likely contribute to the response of the lung following perinatal injury. Together, these factors imply that signals for lung regeneration may be age dependent, emphasizing the need to consider adult and pediatric regeneration as separable processes.

Second, it is possible that the fetal immune system is less well positioned to generate the “correct” level of regenerative signaling in the alveolus after injury, or that the developing alveolus is more susceptible to injury from inflammatory signaling. Our data implies a central role of CXCL and CCL ligands signaling through atypical chemokine receptors in the fetal lung response to LPS; the ACKR family has been implicated in diverse processes in nonimmune cells^62^, especially in directing cancer-associated endothelial biology^84^. Blockade of ACKR receptivity is an area of active anti-cancer drug development^11^, providing possible small molecules which could be considered as chorioamnionitis therapeutics in the future. Taken together, our data emphasize that immune cells are a crucial component of the lung alveolar niche, participating in growth, maintenance, and regeneration of the distal lung throughout life.

### Nonhuman primates bridge a major gap in understanding and treating human disease

Defining actionable pathophysiology underlying complex human diseases remains a challenge, even in the era of unprecedented large data sets from patients. Rhesus is an ideal model organism to address this challenge with translationally focused projects to model human pathophysiology. Our finding that IL1 and TNF blockade protects the lung from injury, and that stromal and innate immune activation elaborates injurious signals which are suppressed with cytokine blockade, provides proof of principle that anti-inflammatory therapies protect the lung from severe injury. These data imply that future therapies targeting the immune system may hold promise for treatment of perinatal inflammation. The quality and flexibility of the Rhesus model opens avenues for biomarker discovery, therapeutic development, and longitudinal analysis for chorioamnionitis and other complex disease. Rhesus also provides a unique future opportunity to directly test assess the causal connections between perinatal inflammation and long-term respiratory health outcomes. Coordination of targeted primate studies with large human disease and organ consortia should be a future priority, as such efforts will enrich our combined knowledge of primate biology, maximize the value of human reference data, and provide novel opportunities for translational biology to develop pathophysiological models and therapies for challenging diseases.

## Acknowledgements

The authors would like to thank the Gene Expression Core (especially Kelly Rangel and Shawn Smith), Confocal Imaging Core (especially director Matt Kofron), and DNA Sequencing Core of the Cincinnati Children’s Research Foundation for extensive technical support. This work has been in extensive collaboration with the California National Primate Research Center, and we would especially like to thank Paul-Michael Sosa, Jennifer Kendrick, and Sarah Lockwood for invaluable help in animal management and care.

## Author Contributions

AT, SS, PK, JG, CMJ, SM, MD, JRS, DB, JK, CC, RR, JSS, and YM gathered and analysed data; DS, YD, KC, JW, YX, NS, EM, MGu, and MGuo assisted with complex data analysis; SGK, IL, LM, CAC, AJ, JAW, HD, and WZ supervised data acquisition and analysis; AT and WZ wrote the first draft of the paper with input from SS and HD; all authors approved the final draft of the manuscript.

## Materials and Methods

### Ethics/Animals

All animal procedures were approved by the Institutional Animal Care and Use Committee at the University of California Davis, and all studies were conducted in accordance with Cincinnati Children’s Hospital Biosafety protocols. Between 2014 and 2019 adult female Rhesus macaques (*Macaca mulatta;* total n=40) were time mated for planned hysterotomies at ~GD 105, ~GD 130 (~80% term gestation), or ~GD150 (see table 1). All animals were obtained within 2d before or after the target gestational age when possible. There were no instances of spontaneous death or preterm labor in the experimental animals. At delivery, fetuses were euthanized with pentobarbital and underwent necropsy immediately. For fetal lungs, the right upper lung lobe was collected for histologic analysis, and the left lung was processed for single cell isolation. Samples spanning different years were assayed and analysed in the same experimental replicates in our laboratory to minimize technical variability.

**Table 1.** Rhesus Sample Information:

| Animal ID | Treatment | GD | Sex | Year | H&E + Imaging | Quantification | Bulk RNA-seq | scRNA-seq |
| --- | --- | --- | --- | --- | --- | --- | --- | --- |
| 33-14 (LMR14-01) | Control (~GD105) | 108 | F | 2014 | ✓ |  | ✓ |  |
| 40-14 (LMR14-02) | Control (~GD105) | 108 | F | 2014 | ✓ |  |  |  |
| 41-14 (LMR14-03) | Control (~GD105) | 107 | M | 2014 | ✓ |  | ✓ |  |
| 24-14 (LMR14-04) | Control (~GD130) | 130 | M | 2014 | ✓ |  | ✓ |  |
| 28-14 (LMR14-05) | Control (~GD130) | 132 | F | 2014 | ✓ |  | ✓ |  |
| 44-14 (LMR14-06) | Control (~GD130) | 136 | F | 2014 | ✓ |  |  |  |
| 209-17 | Control (~GD130) | 135 | F | 2017 | ✓ |  |  |  |
| 423-18 | Control (~GD130) | 132 | M | 2018 | ✓ |  |  |  |
| 427-18 | Control (~GD130) | 131 | M | 2018 | ✓ |  |  |  |
| 430-18 | Control (~GD130) | 130 | M | 2018 | ✓ |  |  |  |
| 432-18 | Control (~GD130) | 129 | M | 2018 | ✓ | ✓ |  |  |
| 506-19 | Control (~GD130) | 129 | M | 2019 | ✓ | ✓ |  | ✓ |
| 515-19 | Control (~GD130) | 131 | F | 2019 | ✓ |  |  | ✓ |
| 529-19 | Control (~GD130) | 130 | M | 2019 | ✓ | ✓ |  |  |
| 32-14 (LMR14-07) | Control (~GD150) | 152 | M | 2014 | ✓ |  | ✓ |  |
| 37-14 (LMR14-08) | Control (~GD150) | 151 | F | 2014 | ✓ |  | ✓ |  |
| 46-14 (LMR14-10) | Control (~GD150) | 144 | M | 2014 | ✓ |  | ✓ |  |
| 218-17 | LPS 16h | 130 | M | 2017 | ✓ | ✓ |  |  |
| 223-17 | LPS 16h | 130 | F | 2017 | ✓ | ✓ |  |  |
| 429-18 | LPS 16h | 132 | F | 2018 | ✓ |  |  |  |
| 436-18 | LPS 16h | 136 | F | 2018 | ✓ |  |  |  |
| 437-18 | LPS 16h | 136 | F | 2018 | ✓ |  |  |  |
| 442-18 | LPS 16h | 130 | M | 2018 | ✓ | ✓ |  |  |
| 512-19 | LPS 16h | 131 | F | 2019 | ✓ |  |  | ✓ |
| 513-19 | LPS 16h | 130 | F | 2019 | ✓ |  |  | ✓ |
| 27-14 | LPS 48h | 128 | F | 2014 | ✓ | ✓ |  |  |
| 36-14 | LPS 48h | 131 | F | 2014 | ✓ | ✓ |  |  |
| 90-15 | LPS 48h | 127 | F | 2015 | ✓ | ✓ |  |  |
| 232-17 | LPS 5d | 133 | M | 2017 | ✓ |  |  |  |
| 233-17 | LPS 5d | 133 | F | 2017 | ✓ | ✓ |  |  |
| 234-17 | LPS 5d | 133 | F | 2017 | ✓ | ✓ |  |  |
| 239-17 | LPS 5d | 134 | M | 2017 | ✓ | ✓ |  |  |
| 240-17 | LPS 5d | 133 | M | 2017 | ✓ |  |  |  |
| 241-17 | LPS 5d | 133 | M | 2017 | ✓ |  |  |  |
| 242-17 | LPS 5d | 131 | F | 2017 | ✓ |  |  |  |
| 500-19 | LPS + anti-TNF + IL-1RA | 135 | M | 2019 | ✓ | ✓ |  |  |
| 501-19 | LPS + anti-TNF + IL-1RA | 133 | M | 2019 | ✓ |  |  |  |
| 509-19 | LPS + anti-TNF + IL-1RA | 135 | M | 2019 | ✓ |  |  | ✓ |
| 510-19 | LPS + anti-TNF + IL-1RA | 133 | F | 2019 | ✓ | ✓ |  | ✓ |
| 511-19 | LPS + anti-TNF + IL-1RA | 131 | M | 2019 | ✓ | ✓ |  | ✓ |
| n = 40 |  |  |  |  | n = 40 | n = 15 | n = 7 | n = 7 |

### LPS and Blockade treatment

Rhesus dams (~GD 130) were randomized to receive 1mL of saline ± LPS (1 mg/mL, Sigma-Aldrich, St. Louis, MO) via ultrasound guided intraamniotic (IA) injection. Hysterotomies were performed 16h, 48h, or 5d post-LPS with delivery of the fetus at GD130 (Table 1). For animals in the combined blockade (Anakinra + Adalimumab) treatment group, the pregnant females simultaneously received both Adalimumab (40 mg SQ 3h prior to LPS and 40 mg IA 1h prior to LPS) and Anakinra (100 mg SQ 3h prior to LPS and 50 mg IA 1h prior to LPS), were given IA LPS as above, and sacrificed 16h post LPS for analysis.

### Histology

The right upper lung lobe was perfused with saline and inflation fixed with 10% formalin at 30 cm H_2_O pressure for at least 24h. Tissue was dehydrated through an ethanol gradient, paraffin-embedded, and sectioned at a thickness of 5μm. Hematoxylin and eosin staining was performed for morphological examination and injury scoring. Following deparaffinization, rehydration, and sodium citrate (10 mM, pH 6.0) antigen retrieval, immunohistochemistry and immunofluorescence was used to detect protein expression on paraffin sections. Antibodies listed in Table 2 were detected using ImmPress HRP Universal antibody polymer detection kit (Horse Anti-mouse/rabbit IgG, Vector Labs, MP-7500), as previously described for mouse in^50^, with TSA plus fluorophores (1:100); antibodies listed in Table 3 were detected using standard immunofluorescence protocols, as previously described in^52^. Sections were stained with DAPI (Invitrogen, D1306, 1:1000) and mounted using Prolong Gold antifade mounting medium (Invitrogen, P36930). TMPRSS2 IHC and FOXJ1 RNAscope were performed as previously reported^42^.

**Table 2.**
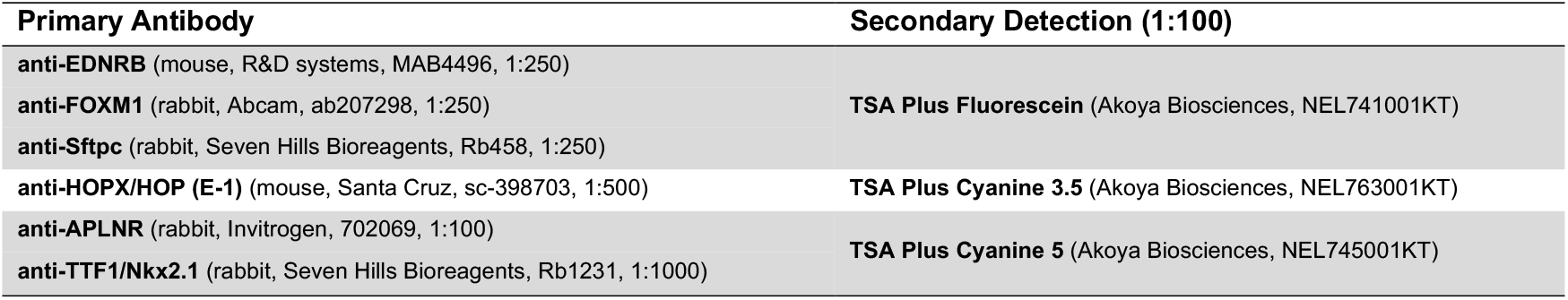
ImmPRESS-HRP + TSA Antibodies and Reagents:

**Table 3.** Standard Immunofluorescence Antibodies:

| Primary Antibody | Secondary Antibody (1:200) |
| --- | --- |
| anti-ABCA3 (guinea pig, Seven Hills Bioreagents, GP985, 1:100) | Goat anti-Guinea Pig IgG Alexa Fluor 555 (Invitrogen, A21435) |
| anti-ACTA2/α-Smooth Muscle Actin (mouse IgG2a, Sigma, A5228, 1:2000) | Donkey anti-Mouse IgG2a Alexa Fluor 633 (Invitrogen, A21136) |
|  | Donkey anti-Mouse IgG Alexa Fluor 647 (Invitrogen, A31571) |
| anti-HOPX/HOP (FL-73) (rabbit, Santa Cruz, SC-30216, 1:200) | Goat anti-Rabbit IgG Alexa Fluor 488 (Invitrogen, A11034) |
| anti-Ki67 (mouse, BD Biosciences, 556003, 1:100) | Goat anti-Mouse IgG1 Alexa Fluor 568 (Invitrogen, A21124) |
| anti-PECAM/CD31 (goat, Santa Cruz, sc-1506, 1:100) | Donkey anti-Goat IgG Alexa Fluor 568 (Invitrogen, A11057) |
| anti-PECAM/CD31 (sheep, R&D Systems, AF806, 1:50) | Donkey anti-Sheep IgG Alexa Fluor 555 (Invitrogen, A21436) |
| anti-pHistone H3 (Ser-10) (rabbit, Santa Cruz, sc-8656-R, 1:100) | Goat anti-Rabbit IgG Alexa Fluor 488 (Invitrogen, A11034) |
|  | Donkey anti-Rabbit IgG Alexa Fluor 555 (Invitrogen, A31572) |
| anti-SCGB1A1/CCSP (rabbit, LS Biosciences, LS-B6822, 1:200) | Goat anti-Rabbit IgG Alexa Fluor 488 (Invitrogen, A11034) |
| anti-TP63 (mouse, Santa Cruz, sc-71827, 1:100) | Goat anti-Mouse IgG Alexa Fluor 488 (Invitrogen, A11001) |
| anti-TTF1/Nkx2.1 (guinea pig, Seven Hills Bioreagents, GP237, 1:200) | Goat anti-Guinea Pig IgG Alexa Fluor 555 (Invitrogen, A21435) |
| anti-TTF1/Nkx2.1 (rabbit, Seven Hills Bioreagents, Rb1231, 1:1000) | Goat anti-Rabbit IgG Alexa Fluor 488 (Invitrogen, A11034) |
|  | Donkey anti-Rabbit IgG Alexa Fluor 488 (Invitrogen, A21206) |
| anti-TUBA1A/Acetylated Tubulin (mouse IgG2b, Sigma, T7451, 1:1000) | Donkey anti-Mouse IgG Alexa Fluor 647 (Invitrogen, A31571) |

**Table 4.**
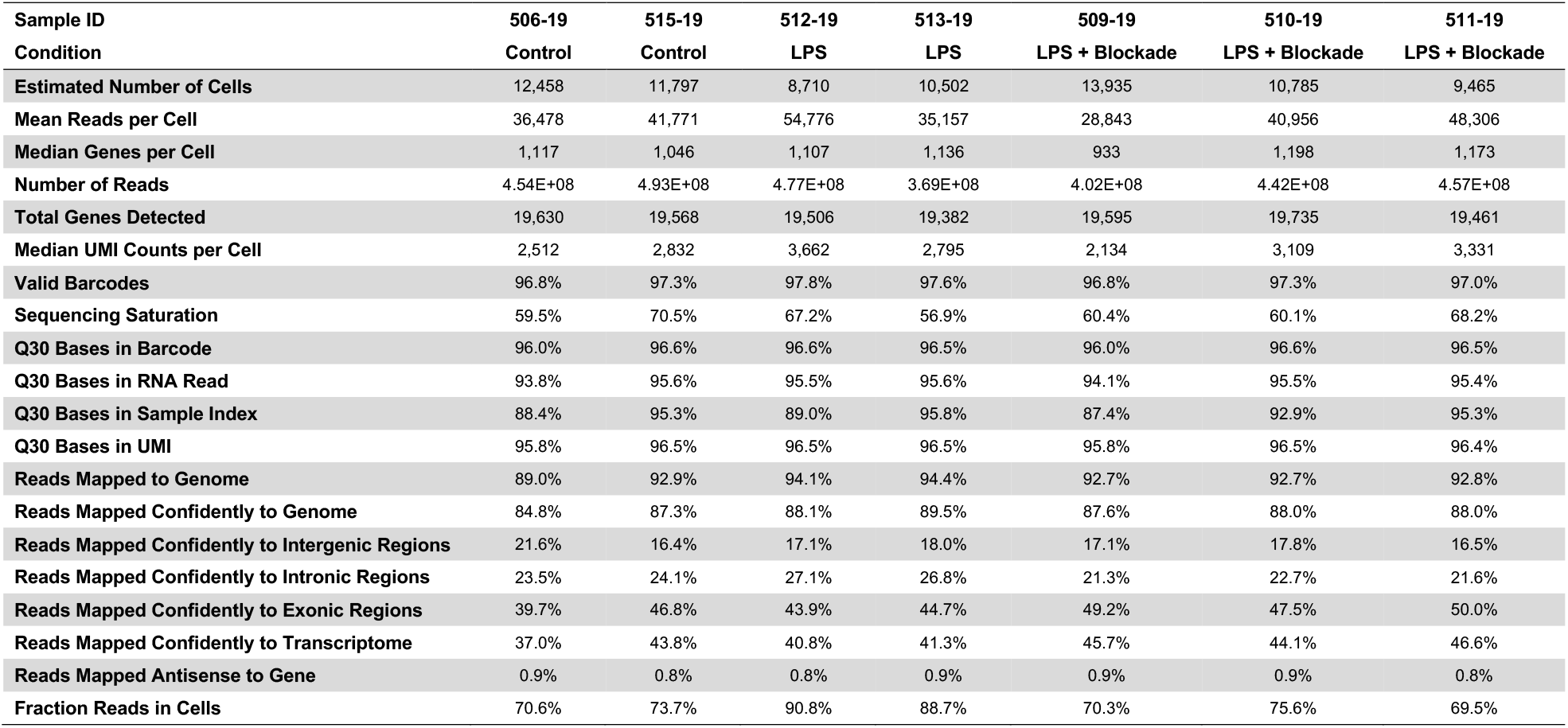
scRNAseq details:

### Imaging

Immunofluorescence and brightfield images were taken using the Nikon Eclipse Ti A1R LUN-V Inverted Confocal Microscope (with 60X Plan Apo IR DIC WI and 100X Plan Apo DIC Oil objectives) and Nikon Eclipse NiE Upright Widefield Microscope (Nikon DS-Fi3 Camera – with Plan Fluor 4x, Plan Apo 10x, Plan Apo VC 20x DIC N2, and Plan Apo 40x DIC M N2, and Plan Apo λ 60x Oil objectives), respectively.

### Image Analysis and Quantification

Representative 40x H&E images from n=3 animals in each experimental group (control GD 130 [27 fields], LPS 16h [37 fields], LPS 48h [39 fields], LPS 5d [31 fields], and LPS + combination blockade [31 fields]) were randomly selected for analysis, including injury scoring and septal counts. Images were deidentified and scored independently (using parameters established by^85^) by two individuals blinded to the treatment condition of the samples. Statistical analyses of these measures were performed using ANOVA with prespecified multiple comparisons in GraphPad Prism 9.0.

### Rhesus lung RNA Isolation and Bulk Sequencing

RNAseq was performed from fetal *Rhesus macaques* at GD105 (n=2), GD130 (n=2) and GD150 (n=3) (see Table 1). Tissue pieces weighing approximately 100 mg were used for RNA extraction. Total RNA was extracted from the snap-frozen tissues using TRIzol Reagent (Life Technologies, 15596-018) and a power tissue-homogenizer (OMNI-TH International, Kennesaw, GA). The RNA was purified using a Qiagen RNeasy Mini kit (Qiagen, 74104) according to the manufacturer’s recommendations. An on-column DNase I digestion step was included in the purification procedure. The concentration of the eluted RNA was determined using a NanoDrop (ND-1000) spectrophotometer. Next generation sequencing of equimolar pools of cDNA libraries was performed by loading a paired-end 75bp rapid flow cell to generate a minimum of 25M raw reads on a HiSeq 2500 sequencing platform (Illumina, San Diego, CA). Raw sequencing data (submitted to Gene Expression Omnibus, GSE178680).

### Rhesus and mouse lung RNA-seq analysis

Partek Flow was used for sequencing alignment and quantification. Ensembl IDs for Rhesus were used for gene annotation. RPKM (Reads Per Kilobase of transcript, per Million mapped reads) was calculated for mRNA abundances analysis. Differentially expressed genes of Rhesus lung development were identified from ANOVA with the corrected p-value (FDR) <0.05 and fold change >1.5 between any age. Differentially expressed genes were subjected to K-mean clustering, genes with two major expression patterns (i.e., induced or suppressed with advancing gestational ages) were selected for functional enrichment analysis using ToppGene^86^. We compared primate data with RNA-seq data from mouse lung development time course data^44^ (GSE122331). Genes differentially expressed during fetal lung development in mouse and Rhesus macaque were directly compared at gene level and key transcription factor level.

### Lung Single Cell Suspension & scRNA Sequencing

After manually dissecting out gross airways, Rhesus fetal lung tissue (~500 mg/tube) was cut into small pieces and transferred into a gentleMACS C-tube (Miltenyi Biotec, 130-093-237) containing 5 mL of digestion buffer (1 mL of Dispase [Corning, 354235], 50 μL of DNase [5 mg/mL, GoldBio, D-301], 100 μL of Collagenase Type I [480U/mL, Gibco, 17100-017] in 9 mL of PBS [Gibco, 10010-023]). C-tubes were processed using the gentleMACS Octo Dissociator with Heaters (Miltenyi Biotec, 130-096-427), running programs “m_lung_01_02” (36 sec) twice, “37C_m_LIDK_1” (36 min 12 sec) once, and “m_lung_01_02” (36 sec) once. Resulting tissue suspension was filtered through 100 μm cell strainer (Greiner Bio-One, 542000) into a 50 mL conical and resuspended in 50 mL PBS. Following centrifugation (1000*g* for 5 min at 4°C) and removal of the supernatant, cells were resuspended and incubated in 5 mL of RBC lysis buffer (Invitrogen, 00-4333-57) for 5 min on ice. Following another centrifugation (1000*g* for 5 min at 4°C) and removal of the supernatant, cells were resuspended in 20 mL of DMEM/F12 (1:1) media (Gibco, 11320-033) (without serum) and filtered through a 40 μm cell strainer (Greiner Bio-One, 542040).Cell counts and viability were determined manually using a hemocytometer. From this suspension, 16,000 cells were loaded into one channel of the Chromium system using the v3 single cell reagent kit (10X Genomics, Pleasanton, CA) by the Cincinnati Children’s Hospital Medical Center Single-Cell Gene Expression Core.

### Alignment & Quality Control

Raw sequencing data (submitted to Gene Expression Omnibus, (GSE169390, reviewer token:qxatqgkyhxyrhkl) were aligned to the Rhesus macaque reference Mmul_10 with Cell Ranger 3.0.2 (10X Genomics), generating expression count matrix files. Cells with fewer than 500 features or greater than 5000 features, as well as cells that contained greater than 25% of reads from mitochondrial genes, were removed. Putative multiplets were removed using DoubletFinder^87^ (version 2.0).

### Data analysis

The Seurat package (version 3.1.0, https://satijalab.org/seurat/)^43^ in R 4.0.2 was used for identification of common cell types across different experimental conditions, differential expression analysis, and most visualizations. After log-normalization, 5000 variable features were identified using the vst method for each sample, and experimental and control animal data was integrated using the *FindIntegrationAnchors* and *IntegrateData* functions with default parameters. Integrated data was scaled, and regression performed for mitochondrial genes, ribosomal genes, and cell cycle state. PCA was performed using the 5,000 most highly variable genes and the first 30 principal components (PCs), followed by *FindNeighbors()* and *FindClusters()* commands to generate UMAP clustering. Epithelial, endothelial, mesenchymal, and immune populations were identified and clustered via expression of CDH1, PECAM1, COL1A1, and PTPRC respectively, and manually annotated for further clustering. Subset objects of each cell lineage were generated for the combined control, LPS, and blockade datasets. Manual annotation of cellular identity was performed by via identification of differentially expressed genes for each cluster using Seurat’s implementation of the Wilcoxon rank-sum test (FindMarkers()) and comparing those markers to known cell type-specific genes from LungMAP^44^. Cell type annotations were consistent across all three treatment conditions and all objects.

These common Seurat objects were used as the basis for additional analytical tools to explore our scRNA data. We used slingshot v2.0 (https://github.com/kstreet13/slingshot)^48^ and condiments(https://github.com/HectorRDB/condiments)^57^ in R 4.1.0 to determine pseudotime lineage trajectories and assess differential lineages across conditions. Slingshot trajectories were generated using the *slingshot()* command with default parameters and specifying only the starting cell population. For condiments analysis, we used the *topologyTest* and *progressionTest* to estimate differential trajectories between conditions and generate p-values. PEP trajectories were identified by Monocle3 (https://cole-trapnell-lab.github.io/monocle3/)^88^. For ligand-receptor analysis, we used CellChat^58^ (https://github.com/sqjin/CellChat) v1.1 using the *SecretedSignaling* subset of the Human CellChatDB, with default parameters. Differential expression heatmaps, GO (gene ontology), pathway (PathwayCommons), and TF (GO-Elite TFTarget database) differentials were generated using cellHarmony^89^, using the cell-to-cluster associations from Seurat as the label file rather than alignment to the reference (--referenceType None), fold > 1.2 and empirical Bayes moderated t-test < 0.05 (FDR corrected). Genes with the opposite pattern of regulation in combination blockade vs. injury as injury vs. control, were considered restored with treatment (same statistical criterion as above). cellHarmony GO-Elite results are displayed using Graphpad Prism 9.0. Visualizations were generated by these tools, with additional use of ggplot2 when needed.

### Statistical Tests

All data met the assumptions of the statistical tests used. Statistical tests used for single-cell analyses are described in relevant section. For comparing the differences between groups, we used either unpaired two-tailed Student’s *t*-test for two groups or ANOVA with prespecified multiple comparison testing for groups of three or more. These statistical analyses were performed in GraphPad Prism 9.0. Details of statistical tests are reported in each figure legend.

---

**Supplemental Figure 1.**
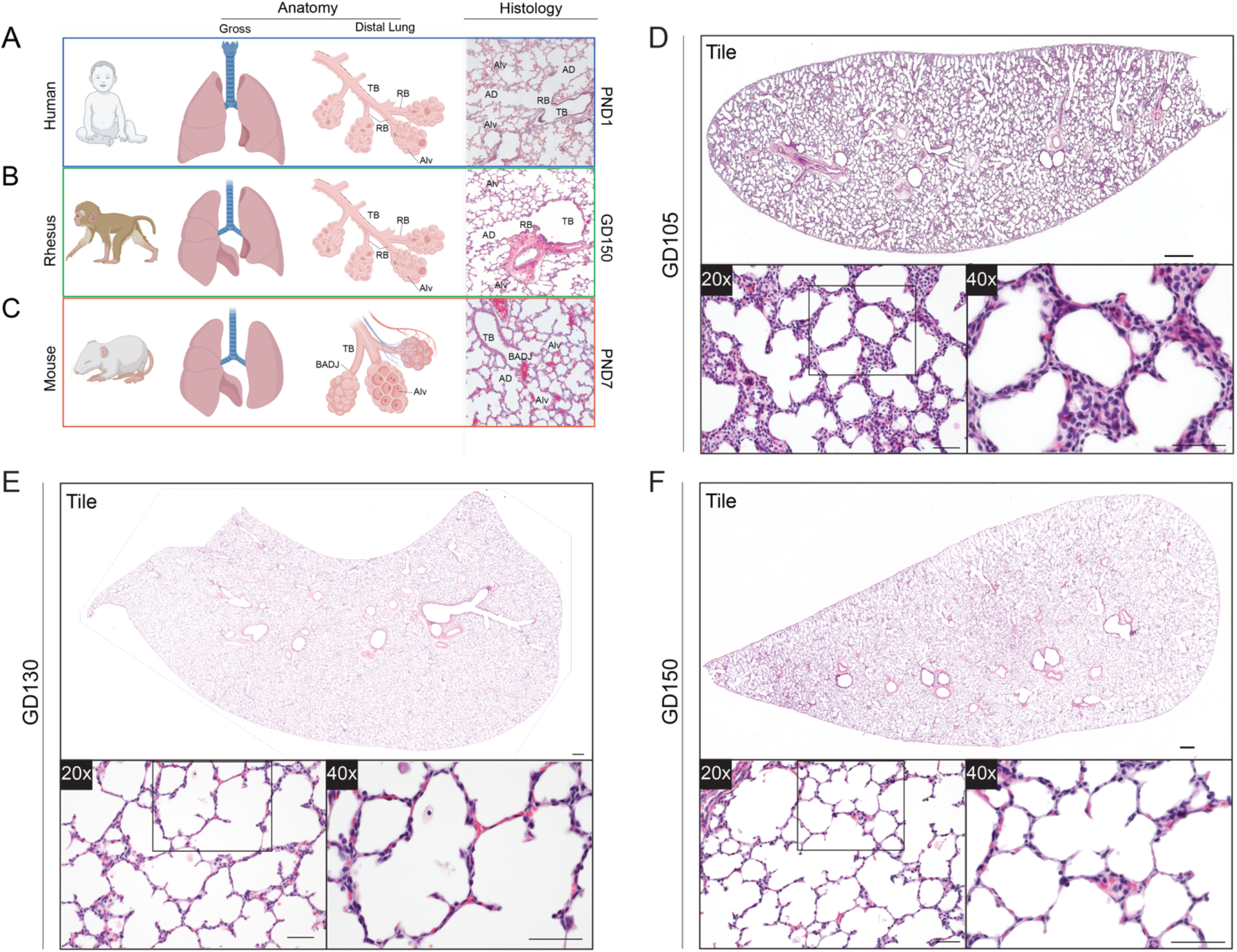
Rhesus macaque lung development models key aspects of human lung organogenesis. A-C) Similarities and differences between human, Rhesus macaque, and mouse lung histologically and anatomically during the alveolar stage of development. Human lung (A) is organized into 3 right lobes and 2 left lobes, with a complex airway tree. The distal terminal bronchi (TB) in humans branch into respiratory bronchioles (RB), which branch into alveolar ducts (AD), which then give rise to multiple alveoli (Alv). Rhesus (B) are similar to human in lobar anatomy, airway complexity, and distal lung anatomy, including respiratory bronchioles, whereas murine lungs (C) are less branched, and terminal bronchi directly give rise to alveolar ducts at a discrete bronchoalveolar duct junction (BADJ). Alveologenesis and alveolar development occur in utero in humans and Rhesus, whereas mice undergo alveologenesis in the first two weeks of postnatal life. (D-F) H&E of Rhesus lung during the canalicular (D; ~GD105), saccular (E; ~GD130), and alveolar (F; ~GD150) stages of third trimester lung development show progressive alveolar maturation and gas exchange development occurs through extension of alveolar septae which are prominent by GD130 (E). By gestational day 150 (F), extensive alveolar structures are present with mature septae. Scale bars = 500 μm [tile scans], 50 μm [20x, 40x panels]. *AD = Alveolar Duct; Alv = Alveolus; BADJ = Bronchoalveolar Duct Junction; GD = Gestational Day; PND = Post-natal Day; RB = Respiratory Bronchiole; TB = Terminal Bronchiole*.

**Supplemental Figure 2.**
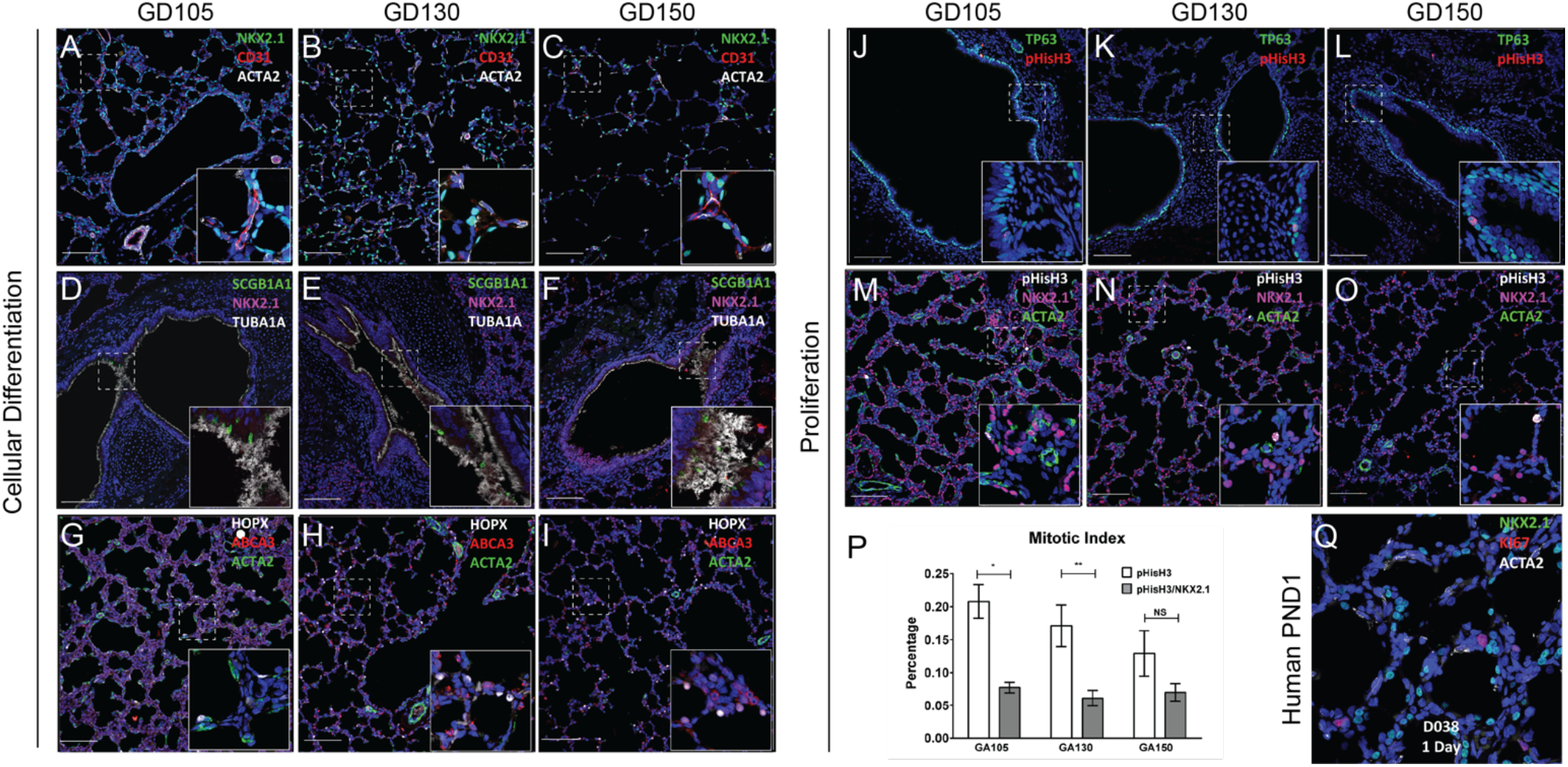
Progressive cellular maturation of the Rhesus lung during third trimester gestation. (A-I) Confocal imaging of epithelial, mesenchymal, and endothelial lineages over the canalicular (A, D, G; ~GD105), saccular (B, E, H; ~GD130), and alveolar (C, F, I; ~GD150) stages of lung development. A-C) Co-development of NKX2.1^+^ epithelium, CD31/PECAM^+^ endothelium, and ACTA2^+^ myofibroblast lineages, with ultimate formation of mature capillary networks in close proximity to both epithelial and mesenchymal lineages. D-E) Progressive differentiation of NKX2.1^+^ epithelium with SCGB1A1^+^ secretory cells and TUBA1A^+^ ciliated cells to develop mature airway structures in preparation for air breathing. G-I) Progressive differentiation and maturation of the alveolar epithelium shown through the presence of HOPX^+^ AT1 cells and ABCA3^+^ AT2 cells forming networks with ACTA2^+^ mesenchymal lineages. J-P) Examination of proliferative dynamics in the developing lung. Proliferation decreases as differentiation increases over the period of alveolarization. Quantification in (P) obtained by counting pHisH3 cells and comparing to DAPI^+^ nuclei (white bars) or NKX2.1^+^ epithelial cells (grey bars). Significantly more proliferation is seen in nonepithelial cells. Q) Comparison of GD150 Rhesus fetal lung to postnatal day 1 (PND1) human lung, showing similar maturation and morphology in the context of both Nkx2.1^+^ epithelial cells and ACTA2^+^ mesenchymal cells. *=p<0.05 and **=p<0.01 by Student’s t-test. Scale bars = 200μm.

**Supplemental Figure 3.**
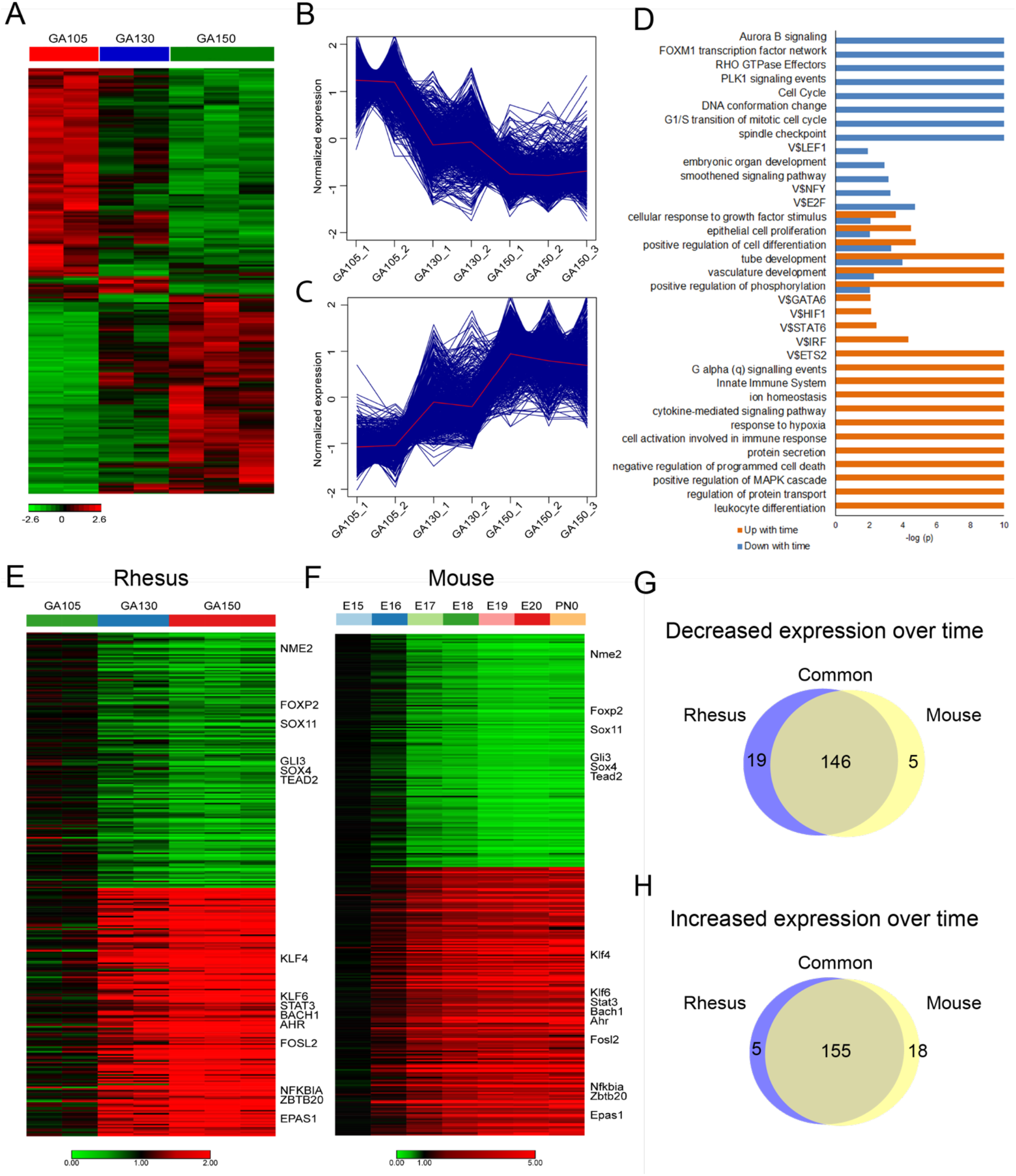
Bulk RNA sequencing of Rhesus lung during development. A-D) Analysis of major differentially regulated genes between GD105 and GD150 in bulk lung RNAseq. Heatmap of differentially regulated genes during Rhesus lung maturation (A) shows significant changes, with distinct gene sets decreasing (B) or increasing (C) across development. GO analysis using ToppGene^86^ of major up (orange) and downregulated (blue) genes demonstrates progressive maturation and reduced proliferation as development proceeds. E-H) Evaluation of major differentially regulated genes as development proceeds comparing later timepoints of Rhesus to GD105 (E) and later timepoints of mouse lung development^44^ to E15. Highly downregulated (G) and upregulated (H) genes were highly concordant between species, implying conservation of major development processes across mammalian evolution.

**Supplemental Figure 4.**
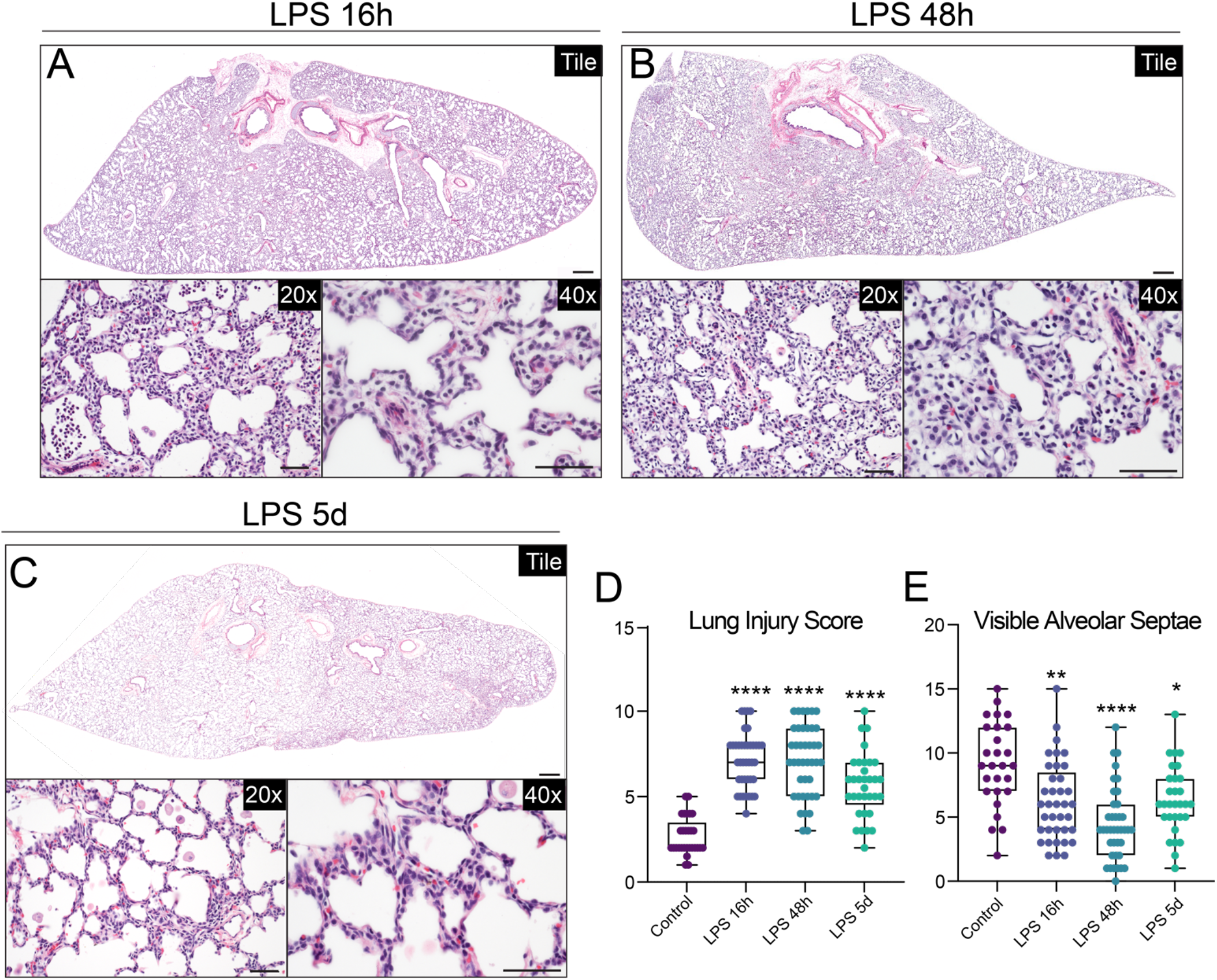
Histological trajectory of LPS-induced inflammatory lung injury. A-C) Histological analysis of LPS-induced lung injury. LPS causes significant alveolar simplification, severe inflammation, and immune cell infiltration, with loss of alveolar septae. Injury progresses throughout 16h, 48h, and 5d post-LPS. (D-E) Blinded image quantification for GA130 controls, LPS 16h, LPS 48h, LPS 5d. (D) Lung injury score, calculated as previously described^85^ (range = 0-10), shows significant lung injury at all time points post-LPS. (E) Counts of visible alveolar septae per field show significant septal loss that is most severe at 16h and 48h post-LPS. * = p <0.05, ** = p <0.01, **** = p < 0.0001 by ANOVA with correction for multiple testing compared to control. Scale bars = 500 μm [tile scans], 50 μm [20x, 40x panels].

**Supplemental Figure 5.**
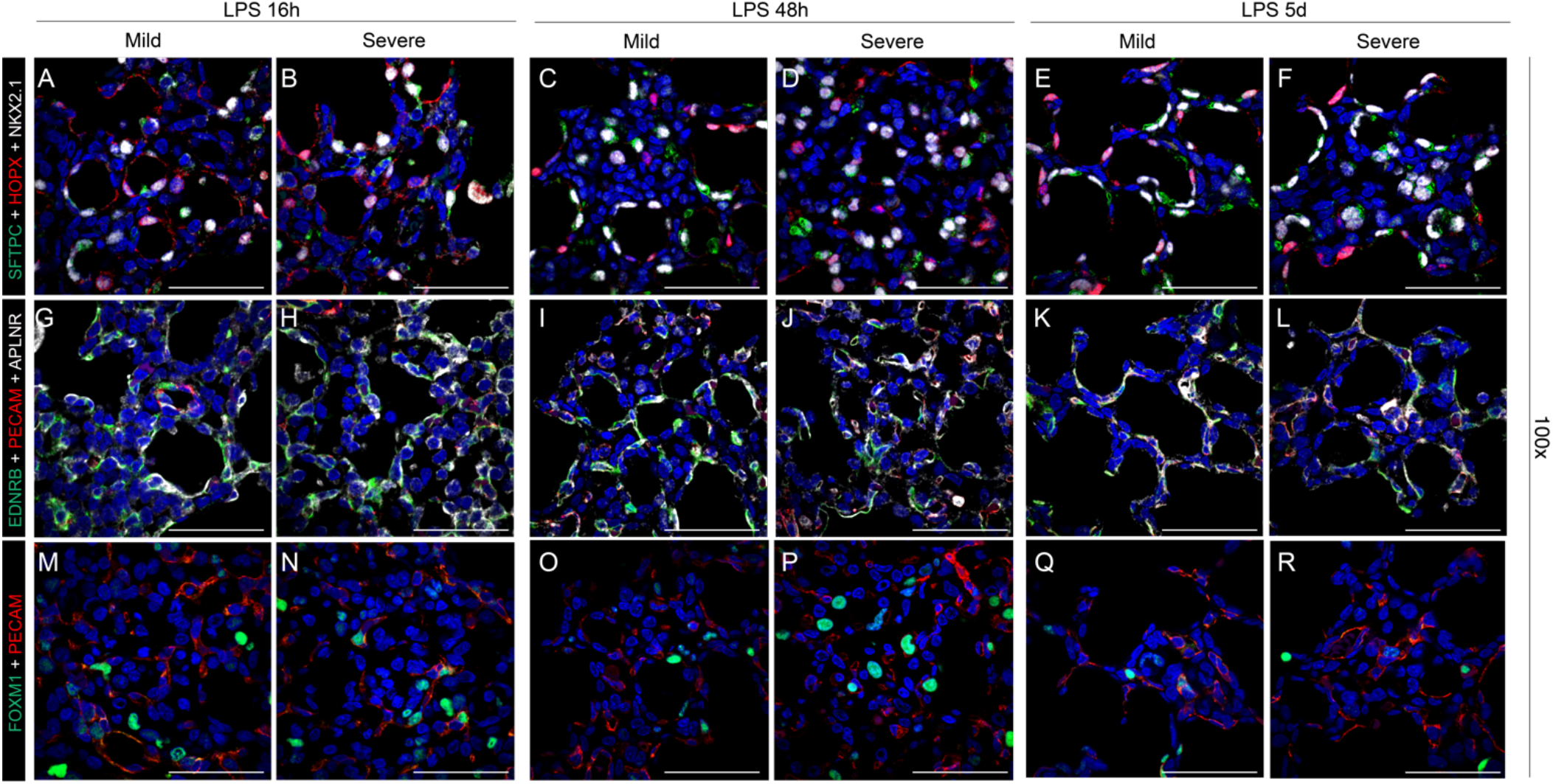
Detailed analysis of trajectory of LPS-induced inflammatory lung injury. Detailed evaluation of epithelial (A-F), endothelial (G-L), and endothelial progenitor (M-R) populations 16h, 48h, and 5d after LPS-induced lung injury. Severe injury observed at 16h and 48h, with mild morphological improvement by 5d post-LPS. All time points show evidence of septal thickening, loss of gas exchange surface area, and disorganization of important cellular populations in the alveolus. (A-F) Destruction of organized alveolar epithelium, specifically loss of SFTPC^+^ AT2 cells (green) and severe damage to/disorganization of HOPX^+^ AT1 cells (red), with the most severe injury noted at 16h and 48h. (L-Q) Loss of EDNRB^+^ alveolar capillaries (green) and disorganization of APLNR^+^ general capillaries (white) following LPS injury. (M-R) Nearly all FOXM1^+^ (green) cells observed are CD31^−^ (red), signifying loss of the CD31^+^/FOXM1^+^ proliferative endothelial progenitor (PEP) at all time points after LPS (compare to figure S7J). Scale bars = 50 μm.

**Supplemental Figure 6.**
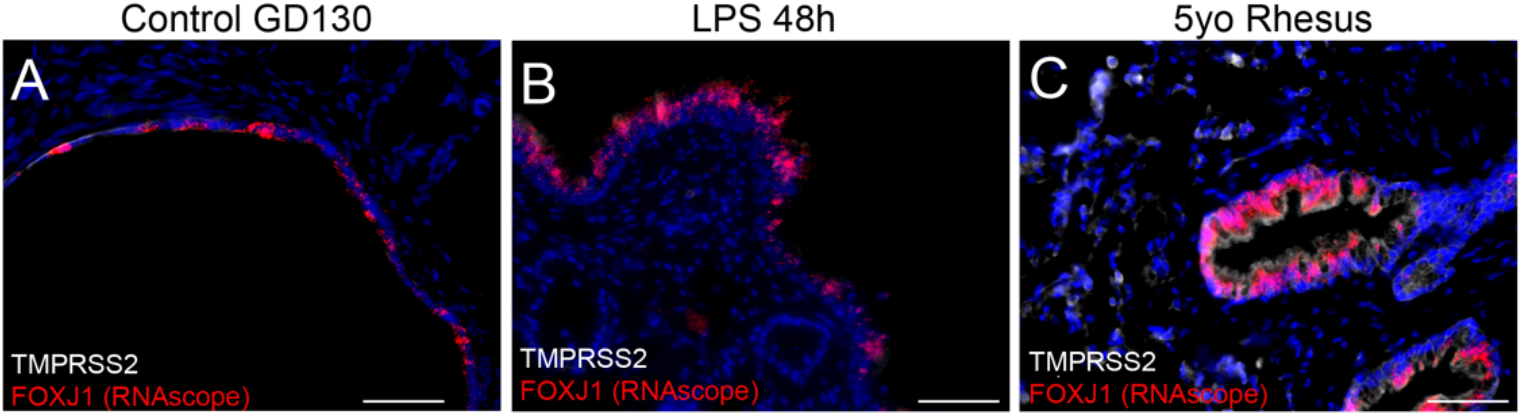
Low level expression of SARS-COV2 protease TMPRSS2 in fetal primate lung. A-C) Evaluation of TMPRSS2 expression in fetal Rhesus lung compared to adult (5yo) Rhesus. Minimal expression of TMPRSS2 is evident in either the airway or alveolar regions of the developing Rhesus lung (A) even following LPS injury (B). Adult Rhesus lung (C), obtained after necropsy for non-pulmonary disease (arthritis), shows significant TMPRSS2 expression in both airway and alveolar cells, concordant with age-specific expression reported in mouse and human lung^42^. Scale bars = 50 μm.

**Supplemental Figure 7.**
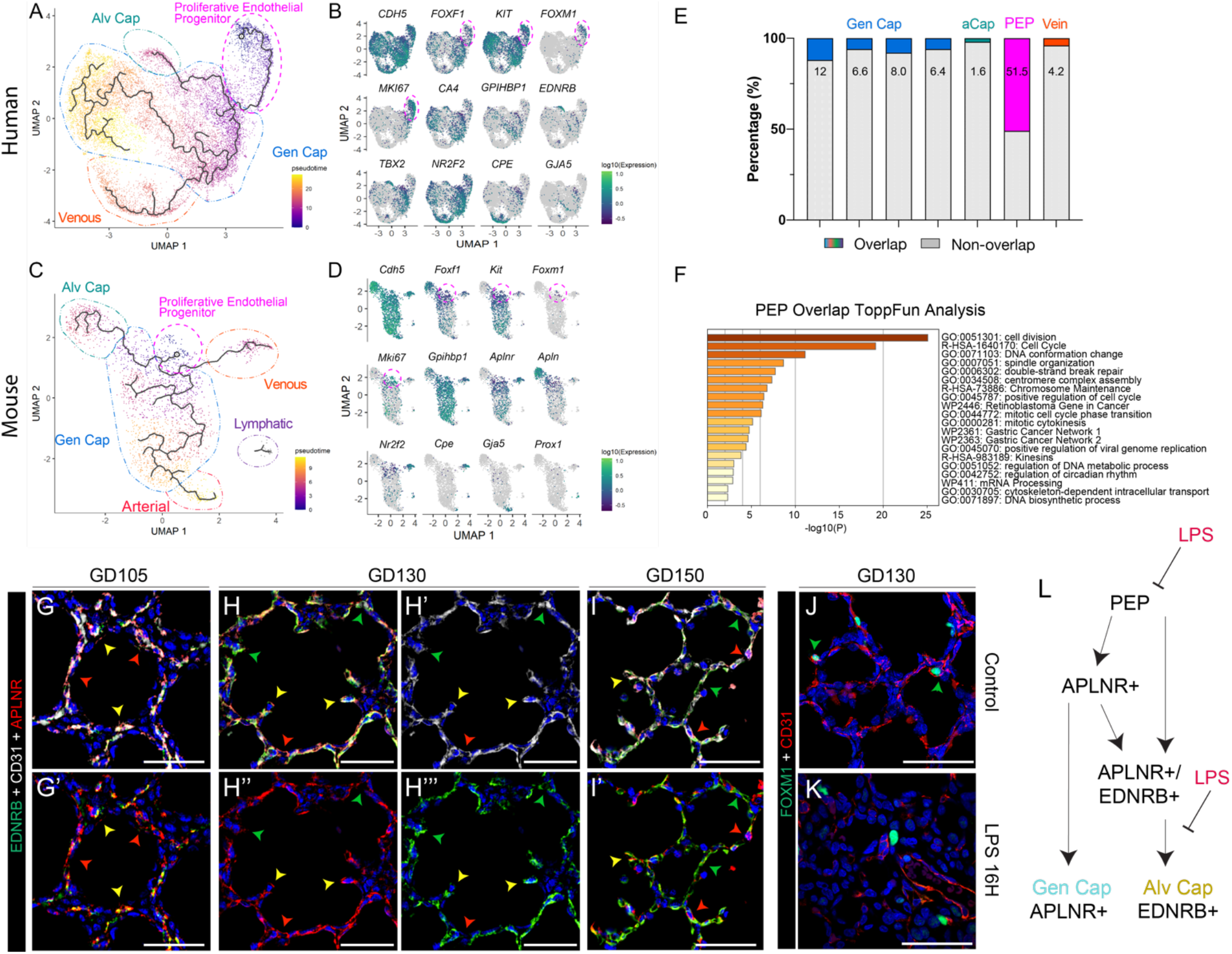
Conservation of the proliferative endothelial progenitor lineage across mammalian evolution. Single-cell RNA-seq (scRNA-seq) analysis identifies proliferative endothelial progenitor (PEP) cells in fetal human lung and developmental mouse lung. A-B) Monocle3^88^ pseudotime analysis of scRNA-seq of vascular endothelial (EC) cells from fetal human lung predicted a differentiation lineage from proliferative EC progenitor to alveolar capillaries, general capillaries, and venous like cells. Data from^54^. Cell identities were defined based on the expression of EC cell subtype markers including pan-EC (*CDH5*), proliferation (*MKI67*, *FOXM1*), EC progenitor (*KIT*, *FOXF1*), general capillary (*GPIHBP1*), alveolar capillary (*TBX2*, *EDNRB*), venous (*NR2F2*, *CPE*), and arterial (*GJA5*) cell markers. Arterial cells make up a small portion of cells in this dataset. The proliferative EC progenitor cell co-expressed *FOXF1*, *KIT*, and *FOXM1*. C-D) The proliferative EC progenitor is also identified in Drop-seq of EC cells from postnatal day 3 and 7 mouse lung^52^; and pseudotime analysis identifies a similar differentiation lineage from the proliferative EC progenitor to alveolar capillaries, general capillaries, venous EC, and arterial EC cells, but not to lymphatic EC cells. Cell identities were defined based on the expression of EC cell subtype markers, including pan-EC (*Cdh5*), proliferation (*Mki67, Foxm1*), EC progenitor (*Kit*, *Foxf1*), general capillary (*Gpihbp1, Aplnr*), alveolar capillary (*Apln*), venous (*Nr2f2*, *Cpe*), and arterial (*Gja5*) cell markers. E) Overlap of top 50 differentially expressed genes from Rhesus PEP with cell populations identified in human lung in (A). F) Enriched terms in ToppGene^86^ GO analysis of PEP genes in overlapping set from (E) G-I) Progressive differentiation of alveolar capillary endothelial cells during alveologenesis, with extensive network at GD150. Red arrowhead = APLNR+ general capillary, green arrowhead = EDNRB+ alveolar capillary, yellow arrowhead = APLNR+/EDNRB+ intermediate endothelial cell. J-K) PEPs are lost in LPS lung injury, with few detectable FOXM1+/CD31+ cells. L) Model of endothelial differentiation, including data from our study. LPS interferes with proper endothelial patterning via loss of PEPs and disruption of alveolar capillary differentiation. Scale bars = 50 μm.

**Supplemental Figure 8.**
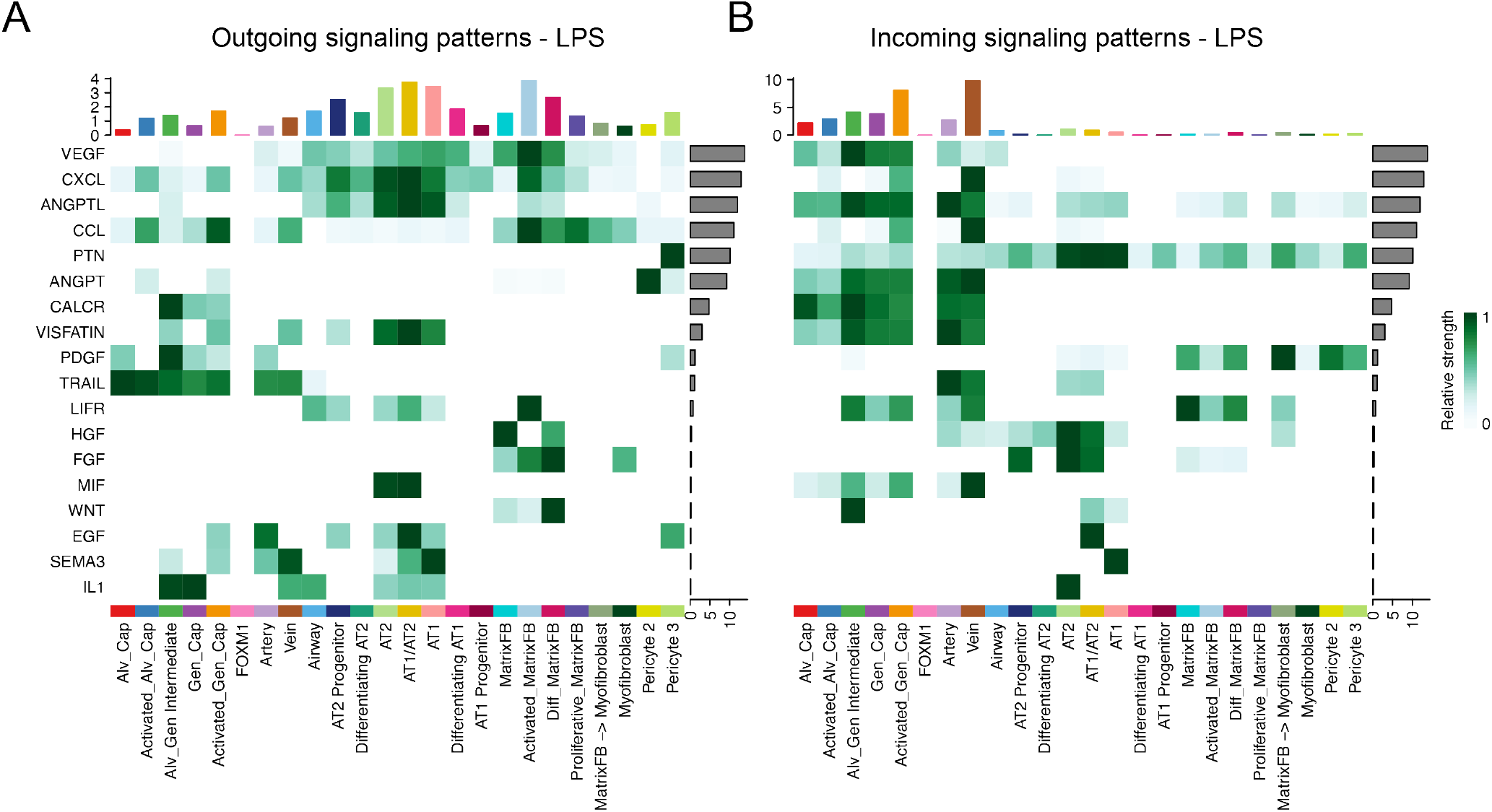
Major signaling patterns in LPS-treated lungs. CellChat analysis of major signaling pathways in milieu of LPS-injured primate lung. See details in Figure 3 legend. A) Outgoing signaling patterns following LPS injury, with significant new inflammatory pathways. Activated matrix fibroblasts contribute to several major inflammatory signaling pathways. B) Incoming signaling pathways following LPS. CCL and CXCL responses are notable in the endothelium and epithelium.

**Supplemental Figure 9.**
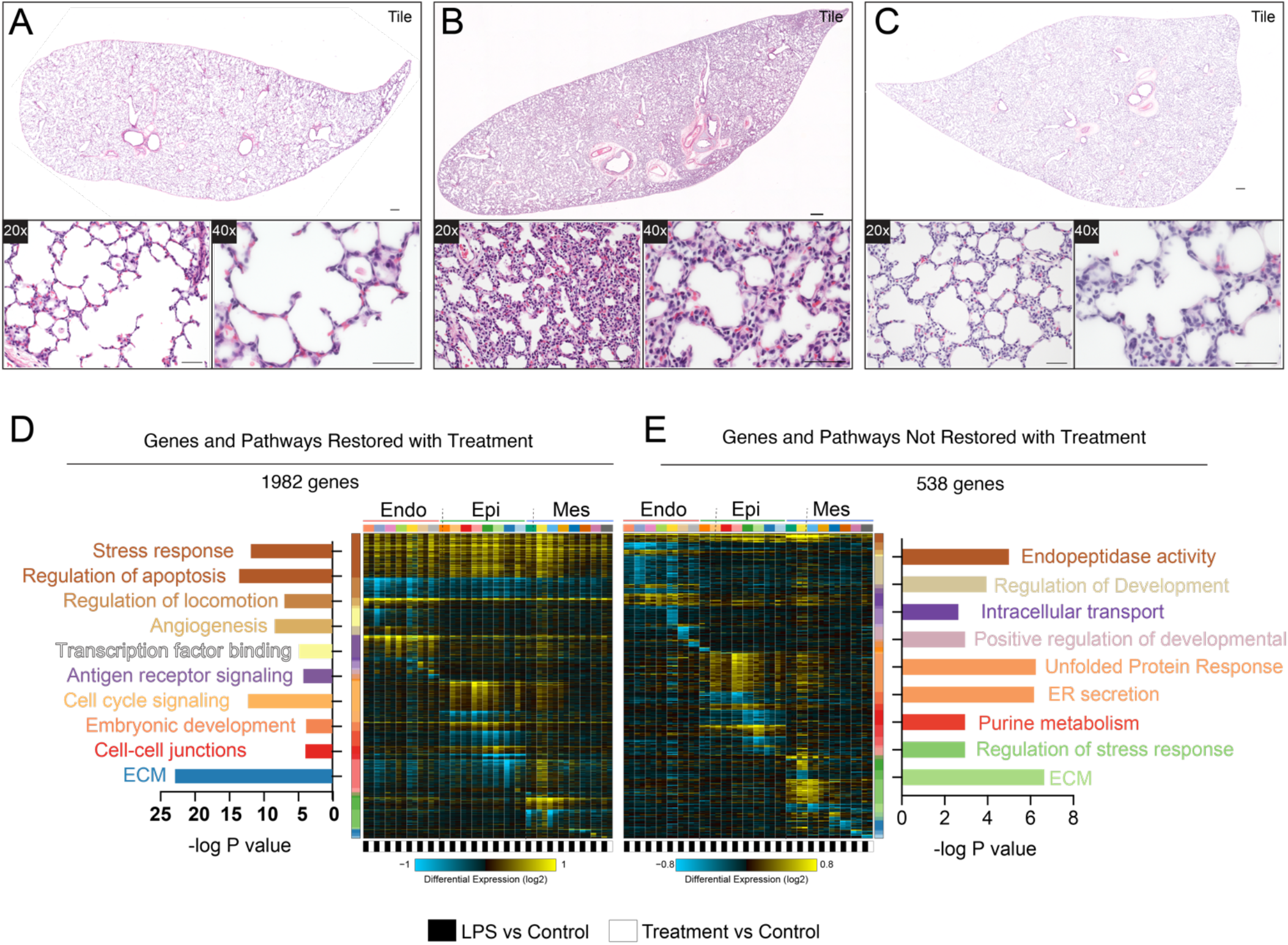
Additional analysis of impact of combination blockade on lung injury from LPS. A-C) Histological analysis of LPS-induced lung injury following combination blockade. LPS causes significant alveolar simplification and severe inflammation, with loss of alveolar septae. Combination blockade protects effectively from this injury. D-E) GO term and pathway analysis of differentially regulated genes by cellHarmony, delineating the effect of combination blockade. Genes and pathways which revert to near baseline expression with blockade are shown in F, while genes and pathways which do not revert are shown in G. Colors of bars are based on differentially expressed gene sets on Y axis. Major stress response pathways, angiogenesis, cell cycle, and antigen response signaling all revert largely to baseline with blockade. Scale bars = 500 μm [tile scans], 50 μm [20x, 40x panels].

